# Mesothelial Plasticity Specifies a Pleural Immune Circuit that Orchestrates Lung Regeneration

**DOI:** 10.64898/2026.06.26.734883

**Authors:** Le Xu, Kevin H Chen, Barsha Dash, Bingpeng Yao, Rongbo Li, Xingan Wang, Zihe Zhou, Vincent Yu, Rebecca Salamon, Pushpinder S Bawa, Minzhe Guo, Zea Borok, Darrell N Kotton, Oliver Eickelberg, Marta Bueno, Xin Sun

**Author notes:** Correspondence (X.S.), (L.X.).

## Abstract

Following injury, predisposition towards regenerative repair and away from degenerative remodeling is central to organismal health, yet the upstream determinants that instruct this fate choice remain poorly understood. Here, we identify mesothelial cell plasticity as a central determinant between regeneration and degeneration by comparing mouse models of pneumonectomy (PNX) versus chronic lung allograft dysfunction (CLAD). While mesothelial cells expand in both settings, following PNX, these cells undergo differentiation through multiple transitional states, culminating in an inflammatory population that orchestrates monocyte recruitment and subsequent tissue regeneration. In contrast, in CLAD, mesothelial cells enrich in an extracellular matrix-enriched state that lacks pro-regenerative signaling capacity. Using single cell epigenomic profiling and in vivo genetics, we define a mesothelium-specific TWIST1-CCL2 axis that governs mesothelium plasticity and signaling, monocyte recruitment and lung regrowth. Together, these findings demonstrate that context-dependent reprogramming of a pleural population, known to protect organs, can be leveraged to drive regeneration.

## Introduction

Adult mammalian organs, including the lung, have long been viewed as tissues with limited regenerative capacity, and maintain homeostasis primarily through regional repair rather than de novo regrowth. This view was challenged by evidence that in human following partial pneumonectomy (PNX), the remaining lobes can undergo substantial compensatory growth, with marked recovery of lung volume and function (Butler et al., 2012; Mizobuchi et al., 2013). In mouse, PNX induces a coordinated regenerative program involving proliferation and differentiation of alveolar type 2 epithelial cells (AT2s), emergence of contractile myofibroblasts for neo-alveologenesis, immune cell recruitment, and vascular remodeling (Chen et al., 2012; Ding et al., 2011; Hsia, 2017; Konerding et al., 2012; Lechner et al., 2017; Li et al., 2020; Liu et al., 2016). Despite these advances, how distinct cellular compartments are spatially and temporally coordinated to rebuild functional alveoli remains incompletely understood.

A defining feature of lung regeneration after PNX is its accelerated growth at the periphery. Mechanical tension is elevated near the pleura of the remaining lobes, where epithelial proliferation and angiogenesis are preferentially activated, suggesting that the pleural compartment constitutes a specialized regenerative niche (Mammoto et al., 2019; Wu et al., 2020). However, the cellular and molecular players that sense and interpret these boundary-associated cues, and how such signals are integrated and relayed to coordinate epithelial, stromal, and immune responses, remain largely unknown.

Another prominent disease condition that features mesothelial remodeling is chronic lung allograft dysfunction (CLAD), the leading cause of long-term mortality after lung transplantation (Sacreas et al., 2019). CLAD is a degenerative disease characterized by persistent and irreversible decline in lung function after transplantation, either as Bronchiolitis Obliterans Syndrome (BOS) or Restrictive Allograft Syndrome (RAS), the latter of which affects as many as 30% of CLAD patients, tends to be more aggressive and is characterized by pleural thickening and parenchymal/(sub)pleural fibrosis (Byrne et al., 2021; Christie et al., 2024; Kurihara et al., 2024; Renaud-Picard et al., 2024; Sacreas *et al*., 2019; Todd et al., 2026; Verleden et al., 2019). While both CLAD and PNX involve surgical manipulation of entire lung lobe(s), CLAD differs from PNX in that it represents a failure of tissue adaptation. The transplanted lung in CLAD succumbs to persistent alloimmune pressure and fails to establish adaptive programs that support tissue homeostasis. In contrast, following PNX, there is no immune rejection response. The remaining lung lobes successfully adapt to altered mechanical forces and physiological demand through coordinated cellular proliferation and differentiation, enabling effective regeneration (Obata et al., 2024). This stark divergence provides a compelling rationale to leverage PNX as a regenerative counterpart to CLAD, enabling us to interrogate whether, and how, mesothelial cell plasticity is differentially deployed to drive adaptive regeneration versus non-adaptive degeneration in the lung.

Mesothelial cells, which connect to form a thin squamous monolayer of the visceral pleura, define the anatomical boundary of internal organs. Similar cells also line the pleural cavity, forming the parietal pleura. Their canonical role is to secrete glycocalyx and provide lubrication to facilitate tissue sliding during respiratory motion (Mutsaers, 2002). Developmentally, lung mesothelial cells arise from mesoderm and contribute to multiple mesenchymal lineages, including smooth muscle cells and fibroblasts (Luna et al., 2026; Morimoto et al., 2010; Que et al., 2008; von Gise et al., 2016). In the adult lung, mesothelial cells are traditionally considered quiescent under homeostatic conditions. However, growing evidence suggests that they retain latent plasticity that can be reactivated upon injury. Studies using a transgenic airway TGFα overexpression mouse model of subpleural fibrosis demonstrated that adult mesothelial cells can proliferate and acquire fibrotic features (Sontake et al., 2018; Sontake et al., 2015). More recently, in a mouse model of lung fibrosis, AAV-based mesothelial lineage-tracing and single-cell transcriptomic data suggest that mesothelial cell-derived progeny cells and extracellular matrix (ECM) proteins promote fibrotic fibroblast formation and fibrosis progression (Fischer et al., 2025; Kadri et al., 2025). These findings implicate mesothelial cells as a driver of pulmonary fibrosis. However, the mechanisms underlying mesothelium function in regenerative and degenerative settings remain poorly defined.

Here, we uncover mesothelial cell plasticity as a signaling center that links boundary-associated cues to tissue outcomes in the adult lung. Through integrated single-cell transcriptomic and epigenomic analyses, lineage tracing, and *in vivo* genetic perturbation, we found that following PNX, mesothelial cells expand and differentiate into multiple transitional populations, ending in a chemokine-producing state that drives immune recruitment and epithelial regeneration. In contrast, in the model of CLAD, while mesothelial cells also expand, they follow a differentiation trajectory that stalls in an ECM- and MHC-enriched state and fail to engage in pro-regenerative signaling. From single cell ATAC-seq (scATAC-seq), we identify TWIST1 as the top candidate driver for mesothelial transition following PNX, and demonstrate *in vivo* that it is essential for mesothelium activation into a signaling center. Furthermore, we show that TWIST1-dependent mesothelium production of CCL2 is required for the recruitment of pro-regenerative monocytes and subsequent alveolar regrowth. These findings support a paradigm in which mesothelial cells act as a context-dependent organizer of the pleural niche, and the range of their plasticity critically instructs whether tissue repair takes on a regenerative versus degenerative path.

## Results

### Multi-compartmental cell reprogramming during lung regeneration

To understand the leading drivers of lung regeneration following PNX, we performed single cell RNA sequencing (scRNA-seq) using either all remaining four lobes or only the accessory lobe which exhibits most regrowth among all lobes at day 7 post PNX and Sham control (Figure 1A and Table S1), when lung volume is substantially increased and AT2 cells (AT2s) are actively proliferating (Liu *et al*., 2016). A total of 21,072 cells (9,744 sham and 11,328 PNX) passed quality control and were used for unsupervised clustering. Cell types from four compartments (epithelium, endothelium, mesenchyme and immune) in the lung were annotated based on established marker genes (Figures 1A and 1B) (Sun et al., 2022). Compared to control, the PNX group exhibits major cellular changes, including 1) the appearance of transitional and proliferative AT2s, 2) the appearance of regenerative myofibroblasts (MFs), and 3) a drastic expansion (∼14-fold) of mesothelial cells (Figures 1C and S1A). Subclustering of epithelial cells revealed that the proliferative AT2s are further separated into two subgroups based on their distinct cell cycle-related gene features: G1/S phase and G2/M phase (Figures S1B and S1C).

**Figure 1.**
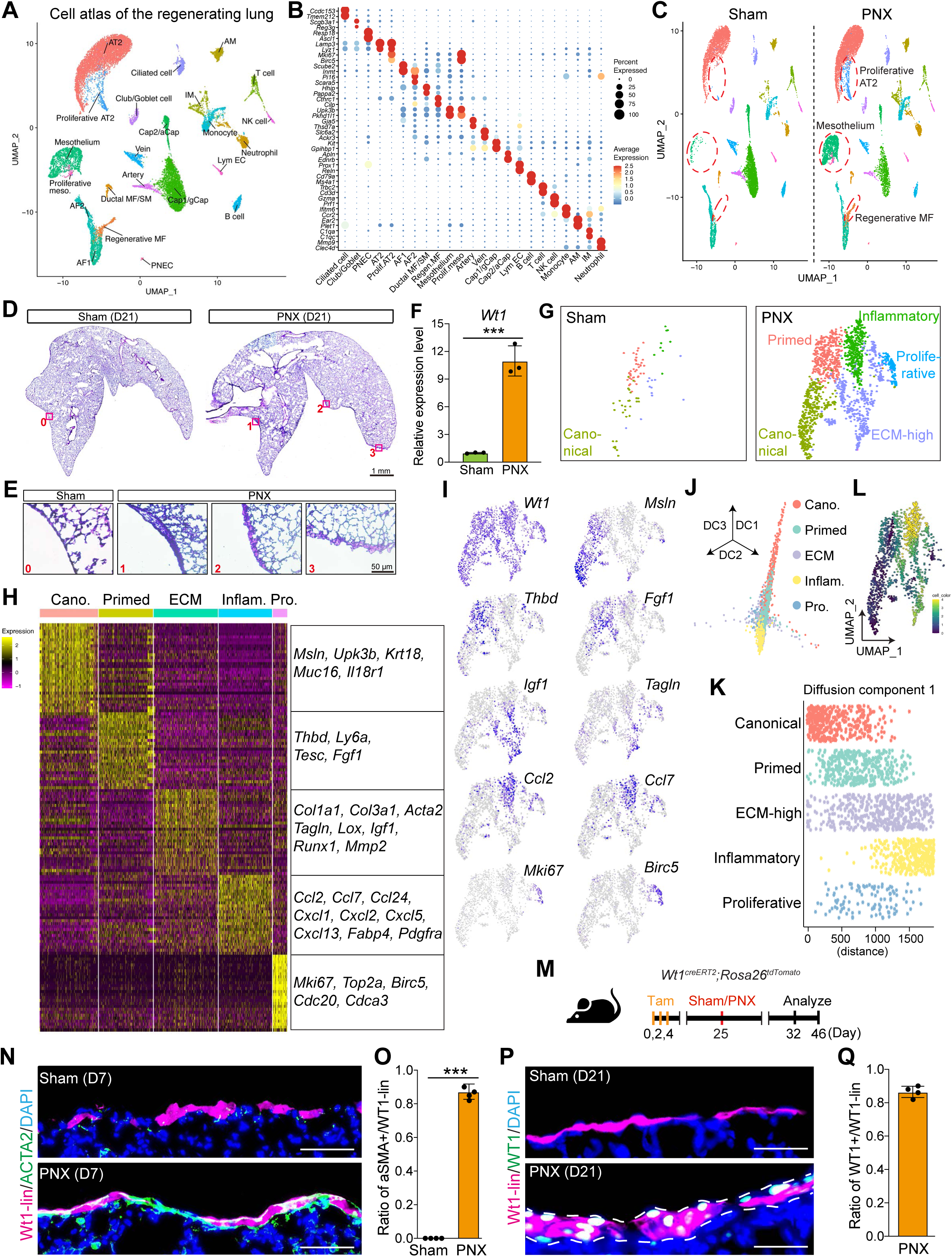
Mesothelial cell reprogramming during lung regeneration. (A) UMAP embedding of integrated cells from sham and PNX lungs at day 7 post surgery. (B) Dot plot profiling of marker genes of each cell type. (C) UMAP embeddings conditioned by sham and PNX. Cell groups with profoundly changed cell numbers were highlighted by dashed red circles. (D-E) Representative H&E staining of sham and PNX lung at day 21 post surgery. The magnified regions at the lung pleura were marked by the red boxes and shown in (E). Scale bars, 1 mm in (D) and 50 µm in (E). (F) Quantification of *Wt1* in sham and PNX lungs at day 7 post surgery, as assayed by RT-qPCR. n=3 for each group. (G) UMAP embeddings of mesothelial cells conditioned by sham and PNX. (H-I) Differentially expressed genes within subtypes of expanded mesothelial cells, as presented by the heatmap. Representative genes were listed in the box at right. Feature plots of selected genes were shown in (I). (J-K) Differentiation trajectory of expanded mesothelial cells, as presented by diffusion mapping of pseudotime. Cells from diffusion component 1 were resolved in (K). (L) Lineage trajectory analysis of expanded mesothelial cells by Monocle 3. (M) Schematic of experimental procedure used for mesothelial cell lineage tracing by *Wt1creER;tdTomato*. (N-O) Representative images of ACTA2 staining with lineage marker tdTomato in sham and PNX lungs at day 7 post surgery. Quantifications of co-detection were shown in (O). n=4 for each group. Scale bars, 20 µm. (P-Q) Representative images of WT1 staining with lineage marker tdTomato in sham and PNX lungs at day 21 post surgery. Quantifications of co-detection in PNX were shown in (Q). n=4. Scale bars, 20 µm. Note that WT1 protein level was below detection in sham lungs, but remarkably increased after PNX.

The appearance of ACTA2+ regenerative MFs in the lung interstitium post PNX has been documented (Chen *et al*., 2012), and we have previously shown that their contraction was required for neo-alveologenesis (Li *et al*., 2020). Yet the transcriptomic features and cellular origin of those regenerative MFs have not been systematically characterized. Subclustering of the mesenchymal compartment revealed that on the UMAP, regenerative MFs are localized adjacent to AF1 cells (Figure S1D), while highly enriched in genes that are typically expressed in MFs, such as *Cthrc1*, *Cemip*, *Col1a1/2, Postn* and *Tagln* (Figures S1E and S1F). Trajectory analysis suggested that these cells originated from AF1 cells (Figure S1G). Consistently, along the differentiation pseudotime, the expression levels of AF1 marker genes gradually decreased, while the expression levels of representative MF genes continuously increased (Figure S1H). To experimentally test if AF1 are able to give rise to regenerative MFs *in vivo*, we generated *Tcf21MerCreMer;tdTomato* mice to trace the AF1/AF2 lineages after PNX (Figure S1I). At day 7 post PNX, we found a clear tdTomato signal in ACTA2+ regenerative MFs (Figure S1J).

The top markers and spatial localization of the regenerative MFs mimicked those of secondary crest myofibroblasts (SCMFs), the well documented cell type that differentiates into ACTA2+ cells at postnatal day (P) 3. Their contraction is required for alveologenesis (Li *et al*., 2020), before undergoing apoptosis at ∼P14 once their roles are accomplished (Hagan et al., 2020). To investigate the transcriptomic similarities and differences between regenerative MFs and SCMFs, we integrated our PNX scRNA-seq data here with our in-house generated scRNA-seq from sorted mesenchymal cells at P8 (Li et al., 2026). On the integrated UMAP, most cells enriched for MF characteristics were clustered together, which were subdivided into SCMFs, Ductal MF/airway smooth muscle (ASM) cells and an intermediate cell population with co-expression of SCMF and Ductal MF-related genes such as *Fgf18*, *Cdh4* and *Egfem1* (Figures S1K and S1M). Most of the cells that constitute these three clusters are primarily derived from P8 lungs. Interestingly, in the adult lung post PNX, most of the regenerative MFs were clustered into the intermediate MF cluster with a minor population remaining with AF1, their cells of origin (Figures S1K and S1L). Differentially expressed gene (DEG) analysis revealed that despite a high degree of profile overlap, regenerative MFs remain distinct from the rest of the developmental MF populations with unique marker genes such as *Cthrc1, Cilp*, *Tnn* and *Inhba* (Figure S1M). To quantify the similarities between regenerative MFs and developmental MFs, enrichment scores of either SCMF-featured or Ductal MF-featured gene set were calculated in each cell group (see Method). Comparative analysis suggests that consistent with their close proximity on the UMAP, regenerative MFs fall in a state between SCMFs and Ductal MF, closer to the signatures of Ductal MF while the P8 intermediate MFs are closer to SCMFs (Figure S1N). Notably, ACTA2 staining post PNX demonstrated that regenerative MFs are abundantly present in the bronchial alveolar duct junction, as well as the interstitial areas in the alveoli (Figure S1O), where, as revealed in thick (99μm) lung section, they form organized networks recapitulating the “fishnet” model reported during postnatal alveologenesis (Figure S1P and Video 1) (Branchfield et al., 2016). Given the extensive increase of gas exchange surface post PNX, findings from our scRNA-seq data lead to the possibility that regenerative MFs in the adult lung are responsible for providing contractile force at either the ductal entrance ring or alveoli for neo-alveologenesis (Figure S1Q). Taken together, our findings here illustrate a concerted array of cross-compartment cellular changes during lung regeneration triggered by PNX.

### Drastic expansion and differentiation of mesothelial cells into diverse subgroups during lung regeneration

To confirm that the expansion of mesothelial cell cluster post PNX reflects a true increase of the population and not a scRNA-seq artifact, we performed histological analysis. We found that the pleura, which is typically composed of a single layer of mesothelial cells, becomes significantly thickened in patches of 3-5 cell layers post PNX (Figures 1D and 1E). Consistently, transcript level of *Wt1*, a gene specifically expressed in mesothelial cells, was increased roughly 10-fold post PNX compared to control (Figure 1F). This increase was also confirmed at the protein level by immunostaining (Figures S1R and S1S). To investigate the cellular complexity within the mesothelial population, we next performed subclustering analysis. Unsupervised clustering yielded 5 subgroups, which were termed canonical, primed, ECM-high, inflammatory and proliferative mesothelial cells based on subgroup-specific marker gene signatures (Figures 1G, 1H and 1I). While *Wt1* is expressed in all clusters, canonical mesothelial cell markers, such as *Msln* and *Upk3b*, are expressed at the highest level in the canonical subgroup, while at a lower level in the primed population and are largely not detected in the rest of the clusters (Figures 1H and 1I). Meanwhile, *Thbd* and *Fgf1* are preferentially increased in expression in the primed population. ECM-associated genes such as *Col1a1, Lox,* and contractility genes such as *Acta2* and *Tagln*, are highly expressed in the ECM-high population. Consistently, trichrome staining confirmed that collagen proteins are elevated in the mesothelial layer after PNX (Figure S1T). For inflammatory population, multiple chemokine genes from CCL or CXCL families are markedly enriched. In addition, a group of proliferative cells was identified by enriched expression of cell cycle genes like *Mki67* and *Birc5*. This proliferative capacity of mesothelial cells was further confirmed *in vivo*, where MKI67 proteins were significantly enriched in the nucleus of mesothelial cells at day 7 post PNX (Figures S1U and S1V).

Diffusion pseudotime and lineage trajectory analysis revealed a strong differentiation trajectory originating from canonical mesothelial cells through primed to ECM-high and inflammatory populations (Figures 1J, 1K and 1L). We note that while both ECM-high and inflammatory populations contain cells with high pseudotime scores, inflammatory mesothelial cells, on average, tend to be located more towards the end of differentiation trajectory. We then used *Wt1creER;R26RtdTomato* to carry out lineage tracing to test the cellular origins of differentiated mesothelial cells *in vivo* (Figure 1M and Video 2). We found that at day 7 post PNX, ACTA2 was ectopically expressed in a significant number of WT1-lineaged cells (Figures 1N and 1O). This finding suggests that the timing when we harvest lungs for single cell study captured the early differentiation of mesothelial cells. Later on at day 21 post PNX, most of the expanded mesothelial cells (∼85%) were traced by the lineage reporter, intermingled with a few WT1+tdTomato-cells (Figures 1P and 1Q). This absence of WT1 lineage marker in WT1+ cells at the lung periphery could either be due to incomplete ability of *Wt1creERT2* to lineage trace all WT1-expressing cells, or it can be due to gain of WT1 expression *de novo* by fibroblasts in the subpleural region (Sontake *et al*., 2018). Consistently, we found a group of *Ebf2*+ AF2 cells in our scRNA-seq that have gained *Wt1* expression post PNX (Figures S1W and S1X). They express *Aldh1a3* as one of the top marker genes (Figures S1E and S1Y), which was also used for annotating subpleural fibroblasts in human lung cell atlas (Sikkema et al., 2023). Taken together, our findings here suggest that adult mesothelial cells post PNX are proliferative and differentiated into heterogenous populations with distinct molecular features.

### Aberrant mesothelial cell reprogramming in chronic lung degeneration

Pleural thickening and fibrosis are well-defined phenotypes in CLAD RAS patients (Byrne *et al*., 2021; Christie *et al*., 2024; Renaud-Picard *et al*., 2024). To address the molecular mechanism, we investigated a CLAD mouse model that recapitulates mixed features of CLAD BOS and RAS, in which the donor lung from genetically engineered mice with one gene mismatch (human *HLA-A2* knock-in) is engrafted onto the recipient lung in *C57BL/6J* background (Smirnova et al., 2019; Smirnova et al., 2022). To compare the underlying mechanism to that of PNX, we performed single cell transcriptomic analysis of this CLAD mouse model (Figure 2A). A full analysis of all cells in the scRNA-seq dataset will be reported elsewhere. In this study, we focus on the mesothelial cells (157 from Syngraft, 546 from CLAD) that are well separated from the rest of the cells in the mesenchymal compartment (Figures S2A and S2B).

**Figure 2.**
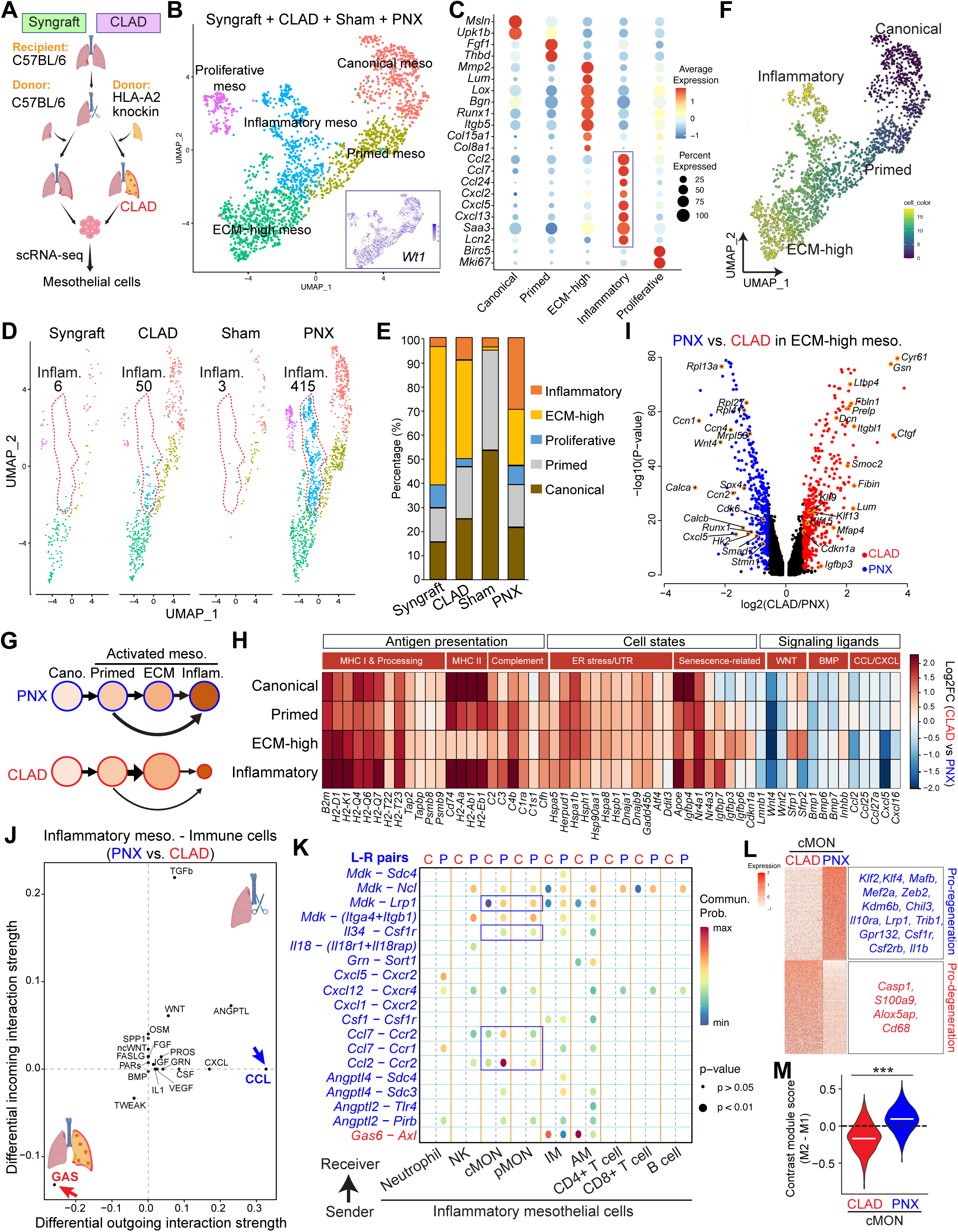
Aberrant mesothelial cell reprogramming in chronic lung degeneration. (A) Schematic of mouse modeling of chronic lung allograft dysfunction (CLAD) used for scRNA-seq. (B) UMAP embedding of integrated cells from syngraft, CLAD, sham and PNX lungs. (C) Dot plot profiling of marker genes of each cell type. (D-E) UMAP embeddings of mesothelial cells separated into different conditions. Normalized proportions of mesothelial cell subtypes under each condition were quantified in (E). (F) Differentiation trajectory of integrated mesothelial cells, as presented by pseudotime analysis from Monocle 3. (G) Diagram presenting the predicted differentiation trajectory of CLAD and PNX mesothelial cells. (H) Representative DEGs identified in at least two subtypes of mesothelial cells between CLAD and PNX, as presented by heatmap. (I) Differentially expressed genes in ECM-high mesothelial cells from CLAD lungs versus PNX lungs, as presented by the volcano plot. Selected genes were highlighted by orange circles and names. (J) Comparative analysis of differential signaling interaction strengths between inflammatory mesothelial cells and immune cells under CLAD versus PNX condition. CCL2 and GAS pathways were highlighted by blue and red arrows, respectively. (K) Top ranked ligand-receptor pairs in the enriched signaling pathways. (L) Differentially expressed genes in *Ccr2*+ canonical monocytes from CLAD lungs versus PNX lungs, as presented by the heatmap. Selected genes were listed in the box at right. (M) Quantification of M1- or M2-biased cell states in CLAD versus PNX lungs, as presented by contrast module scores of M1- and M2-related gene sets (see Method for details).

UMAP clustering showed that similar to PNX, *Wt1*+ mesothelial cells are expanded in the CLAD group compared to Syngraft control (Figures S2C, S2D and S2E). To investigate if similar subpopulations of mesothelial cells were differentiated in CLAD versus PNX, mesothelial cells from each condition (Syngraft, CLAD, Sham and PNX) were integrated for subsequent clustering analysis. Consistent with the integrated PNX and sham control object (Figure 1G), mesothelial cells were categorized into the same 5 subpopulations, each with distinct signatures (Figures 2B and 2C). Interestingly, condition-specific UMAPs showed that, in contrast to other subpopulations that are present in similar proportion in both PNX and CLAD, inflammatory mesothelial cells are preferentially present in PNX and remarkably depleted in CLAD lungs (Figures 2D and 2E). Lineage trajectory analysis demonstrated that, consistent with the lineage hierarchy observed in PNX (Figures 1J, 1K and 1L), the integrated ECM-high population overall shows a reduced differentiation score compared to inflammatory mesothelial cells, suggesting that they may precede inflammatory subgroup along the differentiation trajectory (Figure 2F). However, we cannot rule out the possibility that there is an earlier split in the trajectory from primed to both ECM-high and inflammatory states (Figure 2G). In either of the routes, inflammatory mesothelial population is a prominent target cell state in PNX, while ECM-high is the enriched target cell state in CLAD.

To determine the mechanisms underlying differential mesothelial reprogramming between CLAD and PNX, we performed pairwise differentially expressed genes (DEG) analysis in each of the mesothelial subtypes between CLAD and PNX. We found a total of 1,034 genes upregulated in CLAD and 2,115 genes upregulated in PNX in at least one mesothelial subtype. Among them, 211 (20%) and 273 (13%) genes, respectively, were shared across all subtypes (Figures S2F and S2H). We next categorized DEGs based on their biological functions (Figures S2G and S2I). Interestingly, we found that an extensive panel of MHC class I (*B2m*, classical *H2-D1/H2-K1*, non-classical *H2-Q4/Q6/Q7/T22/T23*) and class II (*Cd74*, *H2-Aa/Ab1/Eb1/Dma*) antigen-presenting genes, as well as the intracellular peptide-processing machinery (*Tap1/2, Tapbp* and *Psmb8/9/10*), are upregulated in at least two of the four CLAD mesothelial subtypes. Accompanying this, multiple complement pathway genes are upregulated as well. Furthermore, genes associated with cellular stress and senescent pathways are found upregulated in more than one subtype of the CLAD mesothelial cells (Figure 2H). These data together suggest that mesothelial cells in CLAD transition into a stress-burdened state, acquire a non-classical antigen-presenting phenotype to engage with alloreactive T cells in the pleural niche. Conversely, ligands and receptors associated with tissue regeneration were preferentially upregulated in PNX mesothelium, including the AT2-supporting WNT ligands *Wnt2* and *Wnt4* and the anti-fibrotic morphogens *Bmp1, Bmp6* and *Bmp7* (Figures 2H and S2I). Last but not the least, chemokine genes from CCL and CXCL families, some of which serve as marker genes of inflammatory mesothelial cells, are also upregulated in the PNX mesothelial cells (Figures 2H and S2I).

Given that the ECM-high population is more enriched in CLAD compared to PNX (Figures 2D and 2E), we focused on the DEGs in this population. Gene ontology analysis of DEGs revealed that a number of ECM genes (*Dcn, Lum, Fbln1* and *Ctgf*), transcription factors (*Klf9, Klf13* and *Klf15*) and canonical cell cycle arrest marker *Cdkn1a* are upregulated in CLAD (Figures 2I, S2G and S2J). On the other hand, translational and post-translational machineries, exemplified by ribosomal subunits (*Rpl13a, Rpl41, Rpl27* and *Mrpl53*), active cell cycle driver *Cdk6* and transcription factors (*Sox4* and *Runx1*) are upregulated in PNX (Figures 2I, S2I and S2K). Taken together, our data suggest that mesothelial cell reprogramming adopts regenerative or degenerative characteristics in a context-dependent manner.

To determine how differential activation of mesothelial cells in PNX vs CLAD could rewire the cellular interactome, especially with immune cells in the regenerative versus degenerative niche, we performed ligand-receptor analysis between inflammatory mesothelial cells and immune cells under PNX versus CLAD (Figures S2L and S2M) (Smirnova *et al*., 2022). We found that CCL family of chemokines rank on top of the outgoing signals with increased strengths in PNX versus CLAD (Figure 2J). Expression enrichment analysis of ligand-receptor pairs suggested that following PNX, within the large CCL family members, CCL2/7-CCR2 axis dominates the interaction flows from inflammatory mesothelial cells to monocytes and NK cells (Figures 2K, S2N, S2O and S2P).

Following recruitment into tissues, monocytes differentiate into a spectrum of myeloid cells that are functionally polarized towards either pro-inflammatory/pro-fibrotic or anti-inflammatory/pro-regenerative characteristics in a microenvironment-dependent manner (Bain and MacDonald, 2022; Chen et al., 2023). We next sought to interrogate the molecular features of classical monocytes (cMON) in PNX versus CLAD. In PNX, cMON showed enrichment of transcriptional regulators and immune modulatory genes associated with cellular differentiation and tissue adaptation, including *Klf2*, *Klf4*, *Mafb*, *Mef2a*, *Zeb2*, *Kdm6b, Csf1r and Csf2rb*, as well as genes linked to anti-inflammatory or reparative myeloid functions such as *Chil3*, *Il10ra*, *Trib1* and *Gpr132* (Figure 2L). *Il1b* in cMON was also found upregulated in the PNX setting, which has been shown to directly promote AT2 stemness (Choi et al., 2020) (Figure 2L). In contrast, cMON in CLAD preferentially expressed genes associated with inflammation-related pathways, including *Casp1*, *S100a9*, and *Alox5ap* (Figure 2L). Consistently, gene module enrichment analysis demonstrated that cMON in PNX were more transcriptionally biased to M2-like pro-regenerative characteristics, in contrast to their M1-like pro-inflammatory/fibrotic counterparts in CLAD (Figure 2M). Connecting these immune cell signatures back to the mesothelium, the signaling pathways enriched in PNX compared to CLAD, such as *Cxcl12-Cxcr4, Il34-Csf1r* and *Mdk-Lrp1,* were known to play critical roles during the differentiation of monocytes into pro-regenerative/anti-inflammatory macrophages (Beider et al., 2014; Foucher et al., 2013; Shi et al., 2024; Zhang et al., 2021) (Figure 2K). Taken together, our findings here suggest that context-dependent mesothelial reprogramming could instruct immune profiles that are closely associated with either regenerative or degenerative outcomes in the lung.

### Epigenomic profiling reveals transcription factors controlling mesothelial differentiation during lung regeneration

To address the drivers for mesothelial cell expansion and differentiation following PNX, we assessed chromatin dynamics at single cell resolution by performing scATAC-seq from sorted mesenchymal cells at day 7 post PNX and control (Figure 3A, also see Method). Consistent with our findings in scRNA-seq, UMAP clustering based on accessible peak regions reveals a remarkable expansion of mesothelial cells in PNX, in addition to the appearance of regenerative MFs in close proximity to AF1 (Figures 3B and 3C). Visualization of sequencing reads on marker genes demonstrates strong cell type-specific chromatin opening signatures (Figures 3D and S3A). Consistent with its uniform RNA expression across mesothelial subclusters (Figure 1I), mesothelial lineage marker *Wt1* maintains comparable chromatin openness in all these subclusters (Figure 3D, red box). Also consistent with the scRNA-seq result, predicted solely based on chromatin accessibilities, we found that mesothelial cells underwent largely the same differentiation trajectory (Figure 3E). To investigate the dynamics of chromatin accessibilities in these subpopulations, we performed differentially accessible chromatin (DAC) analysis, including transcriptional start site (TSS), gene body/intronic regulatory elements and distal enhancer elements. We linked each of those DAC peaks with its closest coding genes as their putative DNA cis-regulatory elements (Figure S3B). We found that there is a strong correlation between DAC peak accessibility and transcriptional changes of its corresponding genes, suggesting that a majority of the epigenomic changes impact transcription. For example, *Msln*, a top marker of mesothelium, possesses the highest openness in the peak region at the TSS site in canonical mesothelial cells, and gradually decreased its openness towards a closed state in the inflammatory mesothelial cells (Figures 3F and 3G). On the contrary, *Cdh2* (also called *N-Cadherin*), an adhesion gene previously linked to epithelial-mesenchymal transition (EMT) that is not expressed in mesothelial cells under normal setting, is noticeably elevated, gaining chromatin openness in TSS and intronic regions along the mesothelial differentiation path to the fullest in the inflammatory population (Figures 3F and 3G). A number of distal DNA regulatory elements were also identified where their chromatin accessibility correlates with linked gene expression. For example, we found a previously uncharacterized DNA element about 23kb downstream of *Igf1* gene, whose DNA accessibility steadily increased from near background level in canonical mesothelial cells to the highest in ECM-high population where *Igf1* was identified as one of the top markers from scRNA-seq (Figures S3C, S3D, 1H and 1I).

**Figure 3.**
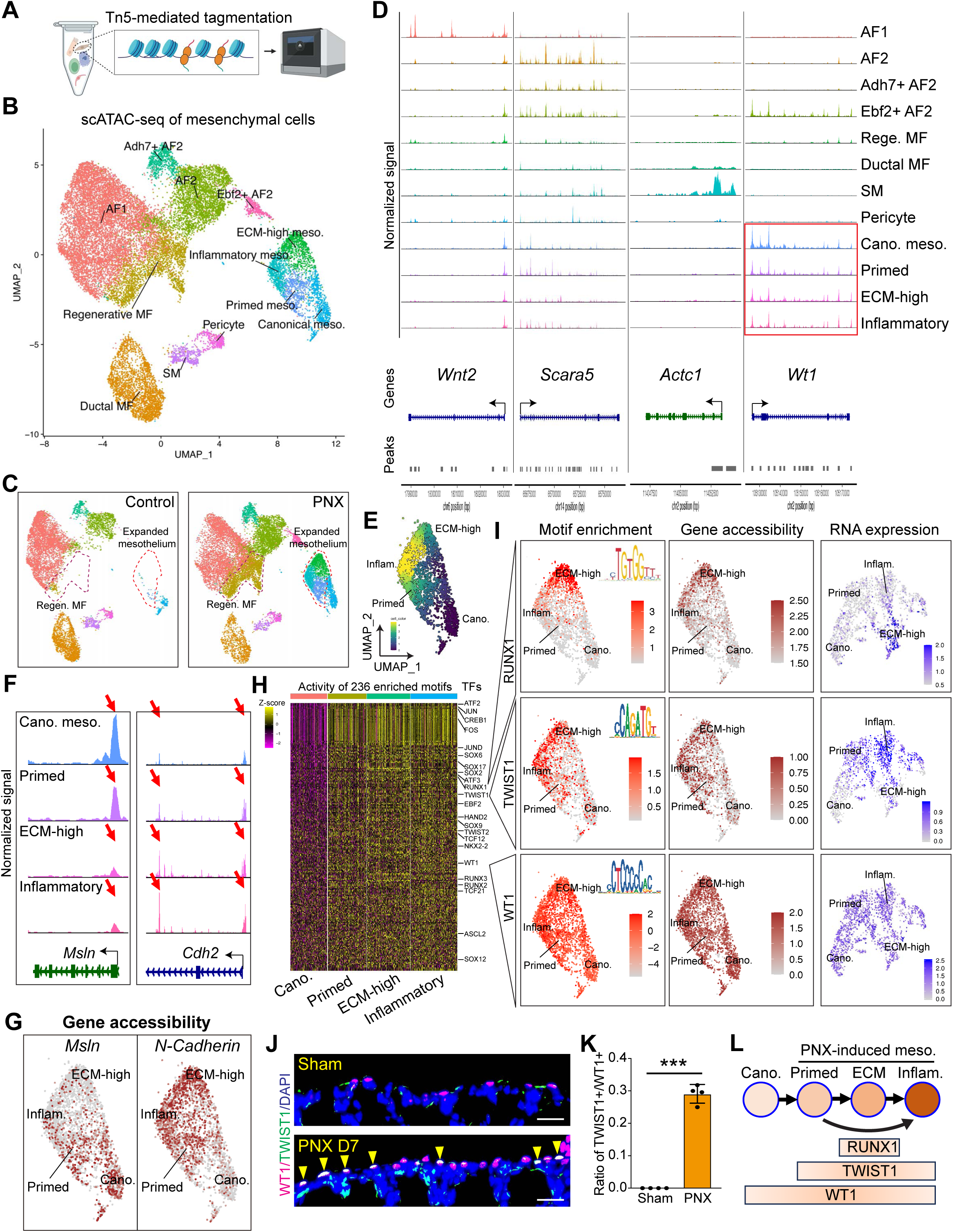
Epigenomic mapping of lung regeneration reveals transcription factors controlling mesothelial differentiation. (A) Schematic of Tn5-based DNA fragmentation and barcoding for scATAC-seq. (B-C) UMAP embedding of integrated mesenchymal cells from control and PNX lungs at day 7 post surgery. Cells from control and PNX lungs were separately shown in (C). Expanded mesothelial cells and regenerative MF are outlined by red dashed lines. (D) Chromatin accessibility of marker genes for selected cell types, as presented by coverage plots. (E) Lineage trajectory analysis of expanded mesothelial cells using scATAC-seq. (F-G) Chromatin accessibility dynamics of *Msln* and *Cdh2* in the mesothelial cell lineage, as presented by coverage plots in (F) and feature plots in (G). (H) Comparative analysis of enrichment scores of TF-binding motifs on open chromatins in canonical versus differentiated mesothelial cells, as presented by heatmap. (I) Feature plots presenting RNA expression (blue), chromatin dynamics (brown) and motif enrichment (red) of selected transcription factors. (J-K) Representative immunofluorescent labeling of TWIST1 on top of WT1 in the sham and PNX lungs at day 7 post surgery. Quantifications were shown in (K). n=4 for each group. Scale bars, 20 µm. (L) Diagram presenting the predicted TF activities within expanded mesothelium.

The direct correlation between gene accessibility and its expression levels in mesothelial lineage enabled us to perform cross-module investigation of DNA binding activities of transcription factors (TF) with their RNA expression features in a cell type- or cell state-specific manner. TF binding motif enrichment analysis in the DNA accessible regions of PNX-induced mesothelial cell states against their canonical counterpart revealed a list of potential TFs that could drive mesothelial cell expansion and differentiation (Figure 3H). Requirement for RNA expression in the given subpopulation narrowed down to the specific candidate genes of interest. As a positive control, we found that mesothelial master regulator WT1 maintained comparable levels of its TF-binding activity, gene accessibility and RNA expression across all mesothelial subpopulations (Figure 3I, lower panel). In comparison, cross-module analysis uncovered TFs more restricted to selected subpopulations, exemplified by the enrichment of RUNX1 activities in ECM-high mesothelial cells (Figure 3I, upper panel). Interestingly, Twist-related protein 1 (TWIST1) acquired *de novo* DNA-binding and transcriptional activity in the primed, ECM-high and inflammatory populations (Figure 3I, middle panel). To independently prioritize candidate TF drivers, we constructed gene regulatory networks (GRNs) for mesothelial populations using our in-house built, PECA2-based pipeline, which jointly models chromatin accessibility and transcriptional output to infer TF-target regulation (Duren et al., 2020; Guo et al., 2023). Consistent with our cross-module analysis, TWIST1 ranked at the top of the predicted TFs (Figures S3E and S3F), reinforcing its candidacy as a key driver of PNX-induced mesothelial differentiation. To confirm the expression change, we performed antibody staining and found that TWIST1 proteins were upregulated in the nucleus of mesothelial cells post PNX (Figures 3J and 3K). Taken together, our findings revealed candidate TF drivers for PNX-induced mesothelial cell expansion and differentiation (Figure 3L).

### TWIST1 is required for mesothelial cell expansion and differentiation

To functionally test the transcriptional drivers of mesothelial expansion and differentiation following PNX, we focused on TWIST1, which is not expressed in canonical mesothelial cells but upregulated across several differentiated lineages, including the inflammatory mesothelium population. TWIST1 is best known for its role in epithelial-mesenchymal transition (Qin et al., 2012). To determine if TWIST1 is required for PNX-induced mesothelial cell reprogramming, we generated *Wt1creER;Twist1f/f;tdTomato* mutant mice (hereafter *Wt1creER;Twist1;tdT*) to inactivate *Twist1* in mesothelial cells while enabling lineage tracing (Figure 4A). At day 21 post PNX, in contrast to the multilayered mesothelial cells traced by reporter in the control lung, *Twist1*-deficient mesothelial cells stayed largely in single layer as they are typically found in the Sham group (Figures 4B and 4C). Consistent with this, the drastic increase of *Wt1* transcript level following PNX in the control lung is much attenuated in the mutant lung (Figure 4D).

**Figure 4.**
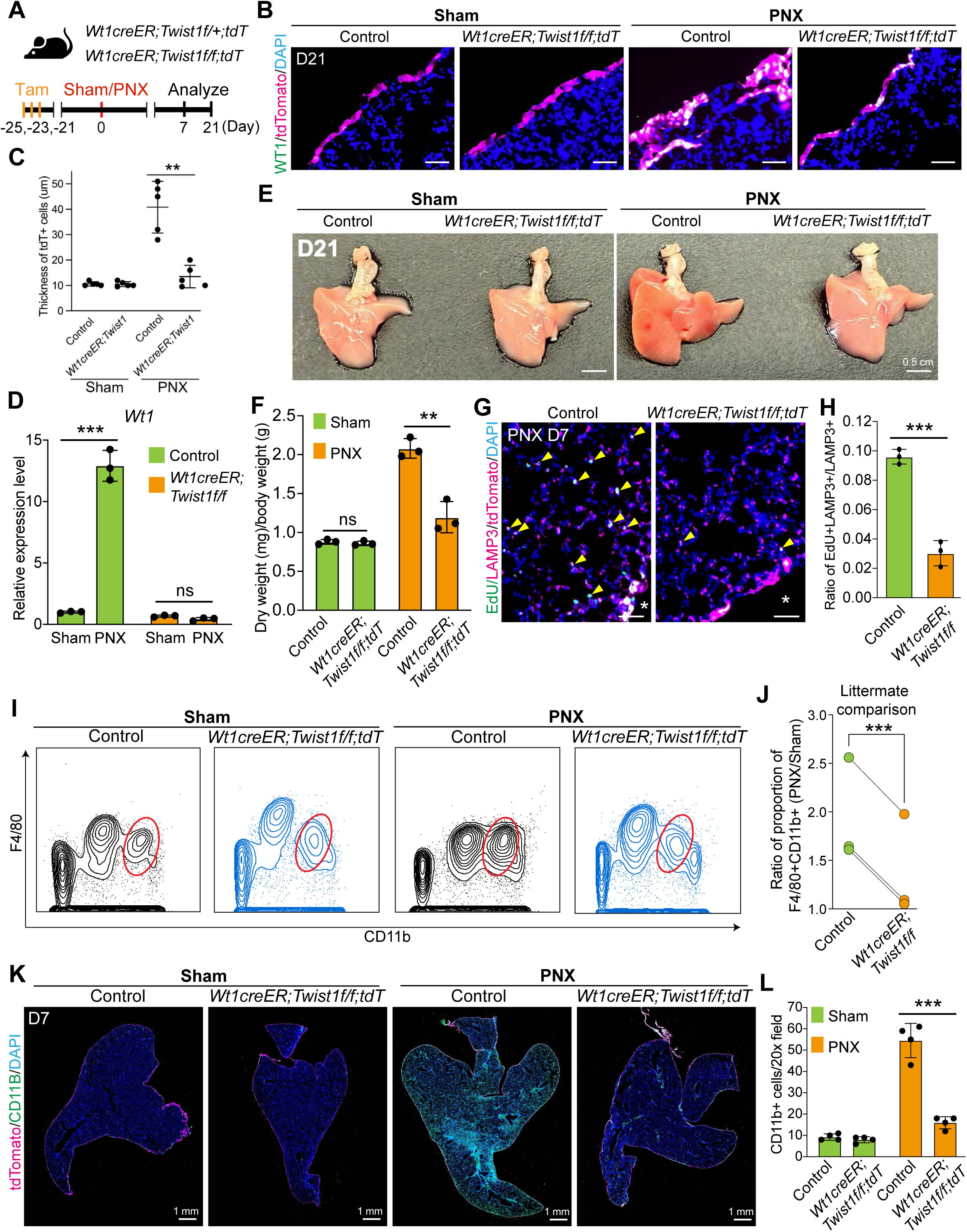
TWIST1 is required for mesothelial cell differentiation. (A) Schematic of experimental procedure used to assay lung regeneration in *Wt1creER;Twist1;tdTomato* mutant mice and corresponding controls. (B-C) Representative immunofluorescent labeling of WT1 on top of lineage marker tdTomato in *Wt1creER;Twist1;tdT* mutant lungs and corresponding controls at day 21 post sham or PNX. Thickness of lineage traced cells was quantified in (C). n=5 for each group. Scale bars, 20 µm. (D) Quantification of *Wt1* in *Wt1creER;Twist1* mutant lungs and corresponding controls at day 7 post surgery, as assayed by RT-qPCR. n=3 for each group. (E-F) Representative images of all right lobes at day 21 post surgery in *Wt1creER;Twist1* mutant lungs and littermate controls. Quantification of dry weight was shown in (F). n=3 for each group. Scale bars, 0.5 cm. (G-H) Representative immunofluorescent labeling of EdU signals on top of LAMP3 in the lung of *Wt1creER;Twist1;tdT* mutants and relative controls at 7 days post PNX. Quantifications were shown in (H). n=3 for each group. Scale bars, 20 µm. (I-J) Flow cytometry analysis of F4/80+CD11b+ monocytes and their derived cells as gated in live CD45+ singlet in the lung of *Wt1creER;Twist1;tdT* mutants and littermate controls at 7 days post surgery. Quantifications within littermates were shown in (J). n=3 for each group. (K-L) Representative immunofluorescent labeling of CD11B with tdTomato reporter in the lung of *Wt1creER;Twist1;tdT* mutants and littermate controls at day 7 post surgery. Quantifications were shown in (L). n=4 for each group. Scale bars, 20 µm.

We next sought to test if the failure of mesothelial cell expansion could hamper lung regeneration. Overall assessment of lung size following compensatory regrowth revealed that the right remaining lobes of *Wt1creER;Twist1;tdT* mutant mice are smaller than those of littermate controls (Figure 4E), and there is a statistically significant reduction in dry weight gain in the mutant lung compared to control post PNX (Figure 4F). As a possible mechanism for this reduction, we found that the ratio of proliferating AT2s, as measured by EdU incorporation at day 7 post PNX, is significantly reduced in the *Twist1* mutant mice compared to controls (Figures 4G, 4H and S4A). Notably, we also found that compared to control lung following PNX, the accumulation of ACTA2+ regenerative MFs in the interstitium, as well as the increase of ACTA2 in the mesothelium, were remarkably diminished in the *Twist1*-deficient lungs (Figures S4B, S4C and S4D). Collectively, these data suggest that TWIST1-dependent mesothelial cell reprogramming is required for lung regeneration.

Recruited CCR2+ monocytes and their derived myeloid cells accumulate coincidently with AT2 proliferation post PNX, and were shown to be required for lung regeneration (Lechner *et al*., 2017). Given the strong chemoattractant features, especially the expression of *Ccl2/Ccl7*, in expanded mesothelial cells, we next sought to test if the recruitment of monocytes and their derived myeloid lineages were disrupted by the absence of expanded mesothelial cells due to loss of *Twist1*. To first determine if TWIST1 could directly regulate transcription of *Ccl2* and *Ccl7*, we examined the presence of TWIST1 binding motif at the accessible peaks around the genic region of those genes. We found that TWIST1 motif sequence was highly enriched in a peak region ∼24kb upstream of *Ccl2* (∼34kb upstream of *Ccl7*), where the chromatin accessibility of this peak was increased from nearly the baseline in canonical mesothelial cells to the highest in the differentiated ones (Figure S4E). Consistent with this chromatin feature, quantification of the RNA expression levels of *Ccl2* and *Ccl7* in control and *Wt1creER;Twist1;tdT* mutant lungs post Sham/PNX revealed that the increase of RNA transcripts of both genes was significantly reversed to close to sham control in the mutant mice post PNX (Figures S4F and S4G). These data suggest that the regulatory role of *Twist1* in the activated mesothelial cells is essential for the upregulation of these chemokines. Consistent with previous findings, flow cytometry analysis demonstrated that CD11b+F4/80+ monocytes and monocyte-derived cells in lungs were robustly increased at day 7 post PNX compared to Sham control (Figures 4I and 4J) (Lechner *et al*., 2017). However, we found that this accumulation was profoundly impaired in the *Wt1creER;Twist1;tdT* mutant lungs (Figures 4I and 4J). Meanwhile, the proportion of CD11b-TCRβ+ T cells was not significantly altered by *Twist1* deficiency following PNX, supporting the specificity of myeloid cell chemotaxis mediated by mesothelial *Twist1* (Figures S4H and S4I). Consistent with our flow data, immunostaining showed that the systemic infiltration of CD11b+ cells post PNX was substantially abolished in the *Twist1* mutant lungs (Figures 4K and 4L). These findings together suggest that mesothelial cell expansion is essential for chemokine upregulation, monocyte recruitment and lung regeneration post PNX.

### Mesothelial cell-derived CCL2 is required for lung regeneration

In the lung, CCL2-CCR2 is a primary and dominant signaling axis for inflammatory monocyte recruitment, disruption of which led to strong reduction of monocytes and their derivates (Kipnis et al., 2003; Lechner et al., 2017), while lung-specific overexpression of CCL2 results in overaccumulation of macrophages at baseline and after injury (Liang et al., 2012). However, the cellular source of CCL2 following PNX remains elusive. To address this, we first analyzed *Ccl2* expression in our integrated scRNA-seq. Feature plot on the global UMAP showed that the mesothelial cells, in addition to the monocytes themselves, are the major cell type expressing *Ccl2* (Figure 5A). Comparing split control versus PNX feature plots, expanded mesothelial cells, but not canonical ones, stand out as an additional source of *Ccl2* following PNX (Figure 5B). Specifically, inflammatory mesothelial subpopulation is where *Ccl2* transcripts were enriched (Figures 1I and 5C). To further validate that CCL2 ligands are expressed in expanded mesothelial cells following PNX, we deployed the *Ccl2-HA-2A-RFP-flox* knockin mice, where in the absence of cre-mediated recombination, RFP is expressed in all *Ccl2*-expressing cells (Shi et al., 2011) (Figure 5D). At day 21 post operation, while rare RFP signals were observed in the Sham control, RFP+ cells were apparent at the lung pleural region post PNX (Figure 5E).

**Figure 5.**
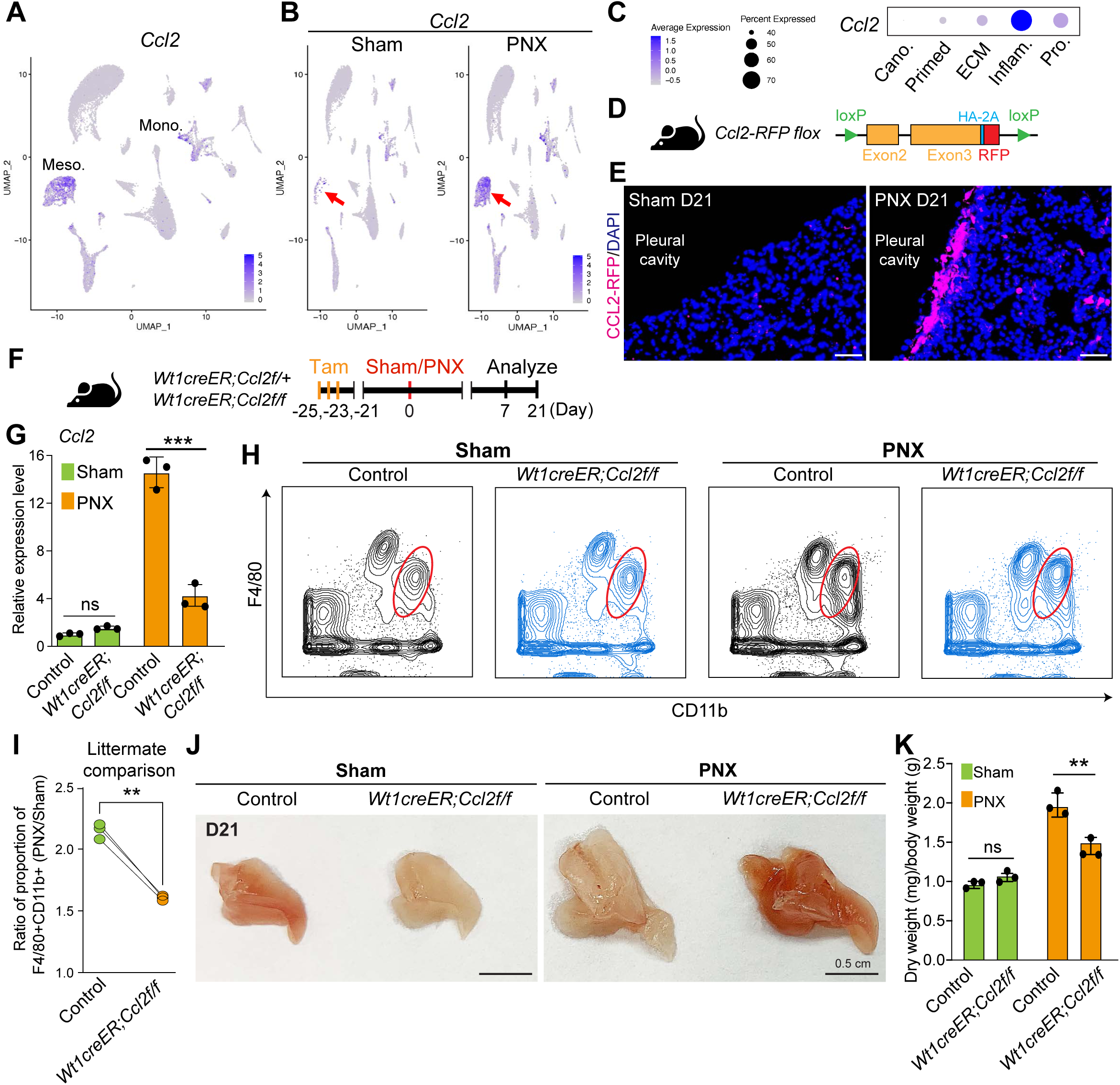
Mesothelial cell-derived CCL2 is required for lung regeneration. (A-B) Birdview of *Ccl2* expression of all cells on the integrated UMAP in (A) and separated conditions in (B). (C) Profile of *Ccl2* expression in expanded mesothelial cells, presented by dot plot. (D) Schematic of *Ccl2-RFP flox* mouse models used to assay endogenous *Ccl2* expression. (E) Representative immunofluorescent labeling of CCL2 at the pleura region in the lung of *Ccl2-RFP flox* at day 7 post surgery. Scale bars, 50 µm. (F) Schematic of experimental procedure used to assay lung regeneration in *Wt1creER;Ccl2f/f;tdT* mutant mice and corresponding controls. (G) Quantification of *Ccl2* in *Wt1creER;Ccl2;tdT* mutant lungs and corresponding controls at day 7 post surgery, as assayed by RT-qPCR. n=3 for each group. (H-I) Flow cytometry analysis of F4/80+CD11b+ monocytes and their derived cells as gated in live CD45+ singlet in the lung of *Wt1creER;Ccl2;tdT* mutants and littermate controls at 7 days post surgery. Quantifications within littermates were shown in (I). n=3 for each group. (J-K) Representative images of all right lobes at day 21 post surgery in *Wt1creER;Ccl2* mutant lungs and littermate controls. Quantification of dry weight were shown in (K). n=3 for each group. Scale bars, 0.5 cm.

To address if *Ccl2* expression in expanded mesothelial cells is required for immune cell recruitment following PNX, we next inactivated *Ccl2* in the mesothelium by crossing the *Ccl2-HA-2A-RFP-flox* mouse line with *Wt1creER* to generate *Wt1creER;Ccl2f/f* mutant mice, and performed PNX together with its littermate controls (Figure 5F). We confirmed that in the mutant, PNX-induced expression of *Ccl2* was substantially inhibited in *Ccl2* mutant lungs compared to controls (Figure 5G). As a consequence, accumulation of monocytes and monocyte-derived myeloid cells post PNX, but not T cells, was significantly impaired by the mesothelial specific loss of *Ccl2* (Figures 5H, 5I, S5A and S5B). These findings suggest that *Ccl2* expression by expanded mesothelial cells is required for monocyte recruitment during lung regeneration.

To determine if the reduction of monocyte lineage caused by loss of *Ccl2* led to defects in lung regeneration, we analyzed the size and dry weight of lung tissues from control and mutant mice following PNX. The overall size of the remaining right lobes is smaller in *Wt1creER;Ccl2* mutant lung compared to control (Figure 5J), with a significant decrease of gained dry weight (Figure 5K). Furthermore, the proliferative capacity of AT2s is also significantly decreased in the mutant lung (Figures S5C and S5D). Taken together, our findings here suggest that the crosstalk between expanded mesothelial cells, specifically the inflammatory mesothelial cells, and monocytes via the CCL2-CCR2 axis is essential for lung regeneration.

## Discussion

In this study, we demonstrate that while the pleural mesothelium is activated in both the PNX model of regeneration and CLAD model of degeneration, only following PNX, it functions as an active regenerative niche for compensatory lung regrowth. Through TWIST1-dependent transcriptional and epigenomic remodeling, mesothelial cells acquire distinct cellular states including an inflammatory state which serves as a signaling center that drives monocyte recruitment and in turn epithelial renewal (Figure 6).

**Figure 6.**
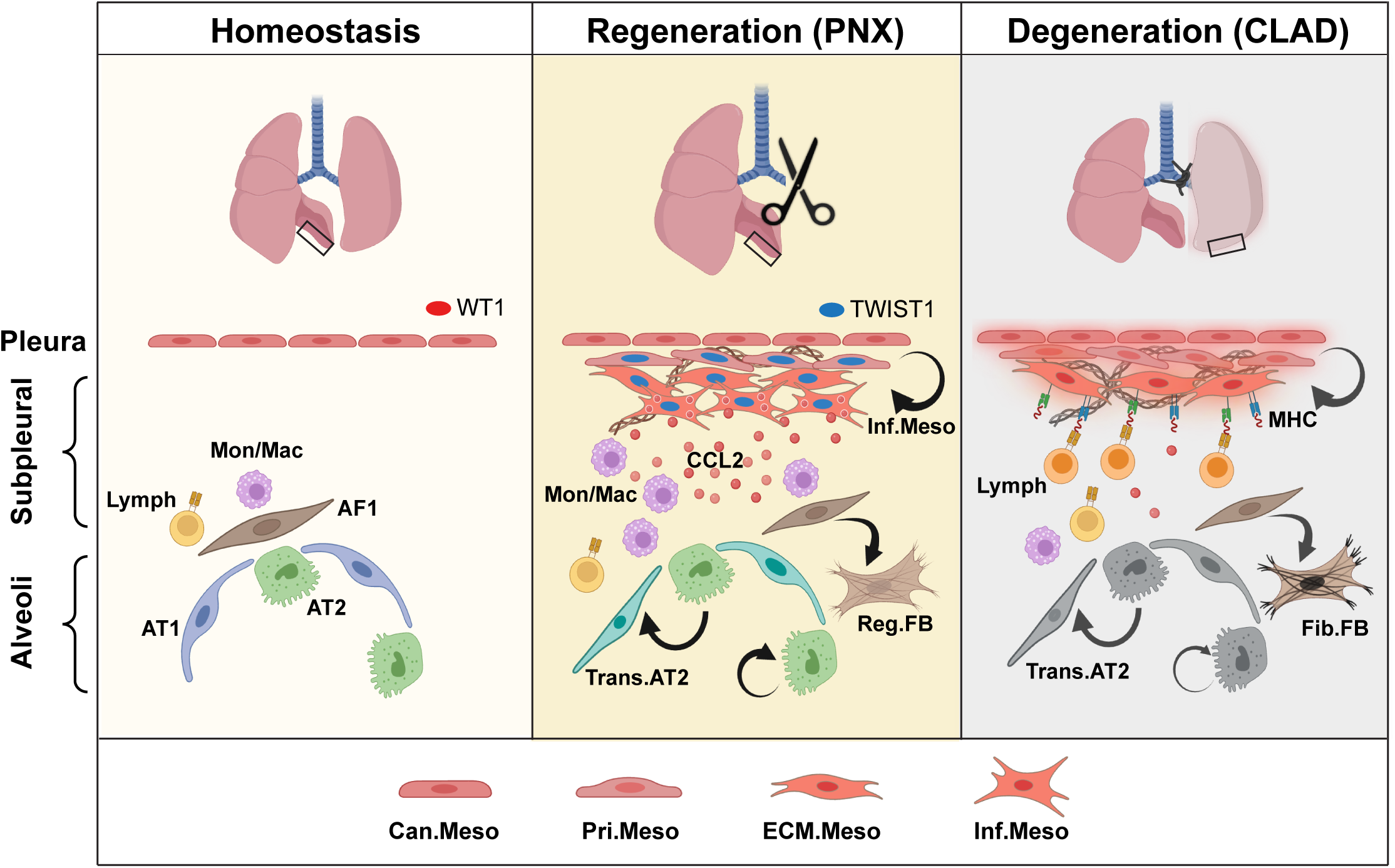
Schematic summary of reprogrammed mesothelial cells function as a signaling hub to orchestrate lung regeneration through recruiting monocytes.

Mesothelial cells share evolutionarily conserved plasticity across organs (Kadri *et al*., 2025). In the injured adult heart, epicardial cells reactivate developmental programs, undergo epithelial–mesenchymal transition (EMT), and give rise to fibroblast- and smooth muscle-like derivatives that contribute to either repair or, in chronic settings, fibrosis (Hesse et al., 2021). Similarly, liver mesothelial cells give rise to fibroblasts following ischemic or inflammatory injury, contributing to fibrosis (Li et al., 2013). These studies establish the mesothelium as a key player in tissue response to injury. However, whether the mesothelium activation triggers adaptive or maladaptive dysplastic repair programs remains a knowledge gap that has profound implication across organs. We show here that with the shared feature of mesothelium expansion in PNX and CLAD, they offer an ideal position for dissection of the instructive roles of the mesothelium in regenerative versus degenerative repair. Our findings reveal that the mesothelial cells take on distinct differentiation paths in PNX versus CLAD, and such difference in plasticity instructs context-specific downstream transcriptional and immune circuits at the organ periphery.

Direct comparison between regenerative PNX and degenerative CLAD mesothelial cell states revealed that the inflammatory mesothelial cells are enriched in PNX and selectively diminished in CLAD. We show that this inflammatory population secretes signals such as CCL2 in a *Twist1*-dependent manner, which in turn is essential to recruit pro-regenerative, anti-inflammatory monocytes, triggering a cascade of repair starting from the pleura. In contrast in CLAD, instead of robust differentiation into the inflammatory state, the mesothelial cells stall and are enriched in the ECM-high state. Beyond this loss of capability to initiate regenerative signaling, CLAD mesothelial cells show a robust upregulation of multiple MHC class I and class II antigen-presentation genes, repositioning the mesothelium not as a passive bystander of allograft injury, but rather as a previously unrecognized non-classical antigen-presenting agent that actively sustains the alloimmune engagement, including T cells and B cells, at the lung pleura and contribute to chronic transplant pathology. This proposed role is consistent with recent findings from single-cell profiling of CLAD lungs that uncovered an activated alloimmune environment populated by distinct immune cell types (Yan et al., 2025), which could be facilitated in part by mesothelial antigen presentation. These findings together suggest that the context of mesothelial reprogramming determines downstream immune behaviors and tissue outcome. Failure to generate the appropriate mesothelial subpopulation may preempt stromal-immune communication, thereby favoring chronic inflammation and progressive degeneration.

A notable parallel emerges when comparing mesothelial reprogramming with lung fibroblast reprogramming during injury. Recent work has shown that in the bleomycin model of lung fibrosis, alveolar fibroblasts (AF1s) give rise to both inflammatory fibroblasts and *Cthrc1*+, ECM-high fibrotic myofibroblasts (Fang et al., 2025; Tsukui et al., 2020; Tsukui et al., 2024). During fibrosis, moving fibroblasts off the inflammatory state and into the ECM-high state is important for limiting alveolar permeability and acute injury. In *Tgfbr2*-KO mice, failed transition into the ECM-high state and prolonged presence of the inflammatory state led to increased leakage of IgM and red blood cells, weight loss and mortality (Tsukui *et al*., 2024). In contrast, our data reveal a different topology for mesothelial differentiation. Here, inflammatory mesothelial cells in PNX are associated with pro-regenerative outcomes, while the ECM-high mesothelium state is enriched in CLAD, a degenerative condition. Thus, while both capable of giving rise to inflammatory and ECM-high states, the specific role of these states in regeneration versus degeneration appears to be distinct. An ability to endow the regenerative characteristics of the mesothelial inflammatory state to alveolar fibroblast inflammatory state could steer the repair program towards regenerative outcomes following fibrotic injury.

Finally, our identification of mesothelial cells as a requisite source of CCL2 after PNX revises an existing model that attributed monocyte recruitment primarily to alveolar epithelial cells (Lechner *et al*., 2017). Given that regenerative growth is spatially biased toward the lung periphery, mesothelium-derived CCL2 provides a compelling mechanism for localized CCR2+ monocyte recruitment at sites of highest mechanical tension. This spatially graded immune influx likely amplifies epithelial proliferation and mesenchymal remodeling in a coordinated manner, reinforcing the boundary as a regenerative active zone. By coupling mechanical cues with chemokine production and immune cell positioning, activated mesothelial cells orchestrate a spatially patterned regenerative program. Together, our findings here posit the lung mesothelial cells as a gatekeeper of immune states and a determinant that could be modulated to achieve effective adult tissue regeneration.

## Methods

### Mice

*Wt1-CreERT2* (010912), *Ccl2-RFP-flox* (016849) and *Rosa26-tdTomato* (Ai14, 007914) mice were purchased from Jackson Laboratory. *Tcf21-MerCreMer* mice were acquired from Dr. Michelle D Tallquist at University of Hawaii. *Twist1-flox* mice were a gift from Dr. Jing Yang at UCSD. Tamoxifen (T5648, Sigma) dissolved in corn oil was administered intraperitoneally at a dose of 50 mg/kg. All mice were housed in facilities accredited by the American Association for Accreditation of Laboratory Animal Care (AAALAC) at University of California San Diego. All animal husbandry and experiments were approved by the Institutional Animal Care and Use Committee (IACUC).

### Pneumonectomy

Procedures were performed as previously described (Li *et al*., 2020). Briefly, 8- to 11-week-old mice were anesthetized with ketamine (100 mg/kg) and xylazine (10 mg/kg) and then intubated and ventilated using a small animal ventilator (Kent Scientific). Buprenorphine (0.1 mg/kg) was used for analgesia during and post operation. The chest wall was incised at the fourth intercostal space, and the left lung lobe was gently lifted out of the chest, ligated at the hilum with string, and completely resected. The incisions on the chest wall and skin were then sutured. Mice were discharged from the ventilator when autonomous breathing recovered. Mice in the sham group underwent the same procedure without removal of the left lobe. For dry weight analysis, right lobes were harvested and wrapped with a piece of foil of known weight. Then it was dried off in the 37-degree chamber for 2 weeks followed by weight measurement. Dry weight of the lungs was calculated by subtracting the foil weight from the final weight.

### Orthotopic mouse transplantation

Mouse lung transplantation was performed as described previously (Smirnova *et al*., 2019). C57BL/6 (syngraft) or HLA-A2 knockin mice (allograft) were used as donors, and C57BL/6 mice were used as recipients. Briefly, donors were anesthetized with the I.P. injection of ketamine/xylazine. The pulmonary artery, bronchus, and vein were carefully separated from one another with blunted forceps, prior to cuffing with 24-, 20-, and 22-gauge cuffs respectively. The recipient mouse was anesthetized with a mixture of medetomidine (1 mg/kg), midazolam (0.05 mg/kg), and fentanyl (0.02 mg/kg); Mice were intubated and connected to a small-animal ventilator (Harvard Apparatus), at a respiratory rate of 120 bpm and a tidal volume of 300 μl. The chest was opened on the left side between ribs 3 and 4, and the native left lung was retracted with a clamp. The hilar structures were carefully separated from one another with blunted forceps. After arrest of the blood and air flow toward the left lung, the cuffed graft pulmonary artery, bronchus, and vein were inserted into the recipient counterparts and ligated with 9-0 sutures. The native left lung was removed, and the incision in the chest was closed with a 6-0 suture, after removing all potential air bubbles from the chest. Antagonist was administered, and the animal was extubated when it showed signs of spontaneous breathing. After the operation, the recipient mice were allowed to recover at 30°C overnight and received buprenorphine for 3 days.

### Tissue preparation and immunofluorescent staining

Mouse lungs were fixed in 4% paraformaldehyde (Electron Microscopy Sciences) diluted in PBS overnight at 4°C. Samples were embedded in either paraffin or OCT (Electron Microscopy Sciences) for sectioning. Antigen retrieval was performed before serum-mediated blocking by using high-pH retrieval buffer (10 mM Tris, 1 mM EDTA, pH 9.0). Primary antibodies with final concentrations used for immunofluorescence staining are: rabbit anti-Wilms Tumor 1 (WT1) monoclonal antibody [10 mg/ml] (ab89901, Abcam), rabbit anti-TWIST1 polyclonal antibody [10 mg/ml] (25465-1-AP, Proteintech), rabbit anti-MKI67 monoclonal antibody [5 mg/ml] (ab15580, Abcam), rabbit anti-N-Cadherin monoclonal antibody [5 mg/ml] (MA5-46719, Invitrogen), Guinea Pig anti-LAMP3 polyclonal antibody [5 mg/ml] (391005, Synaptic Systems), mouse anti-ACTA2 monoclonal antibody [5 mg/ml] (A5228, Sigma) and rat anti-CD11b monoclonal antibody [10 mg/ml] (20-0112, Tonbo). The following secondary antibodies were used with final concentration: Cy3-conjugated goat anti-rabbit IgG [2 mg/ml] (111-165-144, Jackson ImmunoResearch), AF488-conjugated goat anti-mouse IgG [2 mg/ml] (A-11001, Invitrogen), AF488-conjugated goat anti-rabbit IgG [2 mg/ml] (A-11008, Invitrogen) and Cy3-conjugated goat anti-guinea pig IgG [2 mg/ml] (AP108C, Sigma). All images were acquired on the ZEISS AxioImager 2, except for slide scans on Olympus SLIDEVIEW VS200. 20X IF images were used to quantify cells labeled by specific markers. For each condition, at least 3 sections per mouse, and 3 mice per genotype were analyzed.

### Masson’s trichrome staining

Trichrome staining for collagen fibers was performed according to the manufacturer’s protocol (Thomas Scientific, HT15-1KT).

### EdU analysis for cell proliferation

Click-it EdU Cell Proliferation Kit (C10337, Invitrogen) was used to measure the proliferation signal. For EdU analysis in adult mice, 1 ml of 400 mM EdU solution (diluted in PBS, Invitrogen) was intraperitoneally injected. Lungs were harvested 1 hour post injection. Samples were fixed in 4% PFA overnight at 4°C before incubation in 30% sucrose followed by OCT embedding.

### Quantitative PCR (qPCR)

Total RNA from adult lungs was extracted by using Trizol (Invitrogen) and RNeasy Micro RNA extraction kit (Qiagen). Reverse transcription was then carried out to obtain corresponding cDNA using iScript Select cDNA Synthesis Kit (Bio-Rad). qPCR master mix was prepared using SYBR Green reagents (Bio-Rad) and amplification signals were obtained by CFX Connect system (Bio-Rad). At least three biological replicates were assayed for each gene unless otherwise noted. Primers used for qPCR analysis are listed in Table S2.

### Tissue dissociation and sorting of mesenchymal cells

Mouse lungs at day 7 post PNX or Sham were used for cell dissociation followed by cell sorting. Briefly, to remove the circulating blood cells, every mouse was transcardially perfused with 12 ml of cold DPBS (Life Technologies) before harvest. Lungs were then inflated intratracheally with cold lysis buffer (RPMI 1640 (Thermo Scientific) with 10% FBS, 1 mM HEPES (Life Technologies), 1 mM MgCl2 (Life Technologies), 1 mM CaCl2 (Sigma-Aldrich), 0.525 mg/ml collagenase D (Roche), 5 unit/ml Dispase (Stemcell Technologies) and 0.05 mg/ml DNase I (Roche)). Lungs were chopped into the lysis buffer and mechanically homogenized by GentleMACS dissociator (Miltenyi Biotec). Tissues were further incubated in the lysis buffer on the rotator (150 rpm) for 30 min at 37°C, then filtered through a 70 µm MACS SmartStrainer. Red blood cells were removed by ACK lysis buffer (A1049201, Gibco). Cells were further collected by centrifugation at 1500 rpm at 4°C for 5 min (acceleration/deceleration=7), counted with a hemocytometer and then diluted to ∼1X10^6^ cells per ml. After blocking with FcBlock antibody (5 mg/ml) for 10 min at RT, epithelial cells, endothelial cells and immune cells were labeled with biotinylated EpCAM (13-5791-82, Invitrogen, 1:500), CD31 (13-0311-82, eBioscience, 1:500) and CD45 (13-0451-82, eBioscience, 1:500) antibodies respectively, and removed by the magnet-mediated EasySep Cell Isolation technology (19860, STEMCELL). The remaining cells were stained with BV510-CD45 (BioLegend, 103138, 1:100), PE-EpCAM (BioLegend, 118206, 1:100), PE-CD31 (BioLegend, 102408, 1:100) and DAPI (D9542, Sigma). Mesenchymal cells were sorted as DAPI−;EpCAM−;CD31−;CD45− on a FACSAria II high speed sorter (BD Biosciences).

### Single-cell RNA-seq and data analysis

Following FACS, single cell libraries were generated from sorted cells by using Chromium Single Cell 3’ v3 kit (10X Genomics). Sequencing was carried out on the NovaSeq (Illumina) platform at IGM, UCSD. For data analysis, Cell Ranger (version 3.0.2) was used to initially align the raw reads onto the mouse reference genome (mm10) and generate the feature-barcode matrix. Next, R package Seurat (version 4.0) (Hao et al., 2021) was used to perform data quality control, normalization, scaling, principal components analysis (PCA) and the canonical correlation analysis (CCA) based data integration. Briefly, to filter out low-quality cells or doublets, cells with fewer than 200 or more than 6,000 unique features, or more than 15% mitochondrial contents were removed from further analysis. Then “LogNormalize” was used to normalize the feature expression. A total of 2,000 top variable features were identified by function FindVariableFeatures and kept for PCA. Top 20 significant components were chosen to conduct dimensional reduction by uniform manifold approximation and projection (UMAP) with default parameters. Module score analysis was performed by AddModuleScore() in Seurat using following gene panels: 1) SCMF: *Htra1, Nes, Cd248, Car2, Agt, Egfem1, P2ry14, Stc1, Aard, Basp1, Plk2, Pdlim3, Eng, Atp2b1, Nrep, Prag1, Fam129a, Tenm4, Wnt11, Pde5a, 6330403K07Rik, Sema6d, Tmem178, Ptma, Lef1, Rerg, Ckb, Etv1, Myh11* and *Cfl1*. 2) Ductal MF: *Hhip, Scx, Lrrc15, Kcnma1, Sbspon, Gja1, Aspn, Mpped2, Enpp2, Lgr6, Cnn1, Cdh4, Fhod3, Myocd, Grem1, Ssc5d, Lmod1, Pde1a, Map1b, Kcnj8, Synpo2, Net1, Sema3c, Ednrb, Col12a1, Pde10a, Enpp1, Actg2, Mustn1* and *Ptgis*. 3) mesothelial cell: *Wt1, Gpm6a, Upk3b, Krt19, Bicd1, Rspo1, Ccdc80* and *Nkain4*. 4) M1 macrophage: *Tnf, Il6, Nos2, Cxcl9, Cxcl10, Ptgs2, Il12b, Cd80, Cd86, Cybb, Socs1, Ido1, Stat1, Stat5a, Irf1, Irf3* and *Irf5*. 5) M2 macrophage: *Arg1, Mrc1, Cd163, Tgfb1, Chil3, Klf4, Il10, Ccl17, Ccl22, Pparg, Retnla, Stat3, Stat6, Irf4* and *Kdm6b*. The expression features of marker genes were profiled and visualized by R package ggplot2 (v.3.5.1). Lineage trajectory analysis was performed by Destiny (Angerer et al., 2016) or Monocle 3 (Trapnell et al., 2014).

### scATAC-seq analysis

After cell dissociation and sorting as mentioned above, mesenchymal cells were subjected to scATAC-seq following the standard protocol by the Center for Epigenomics at UCSD (Wang et al., 2020). For data analysis, read alignment (mouse genome mm10) and cell barcode demultiplexing were conducted using 10x Genomics Cell Ranger ATAC (v.2.1.0) with default settings. Quality control, transcriptional start site enrichment, data normalization, dimensional reduction and UMAP-based cell clustering were performed using Signac (Stuart et al., 2021). Analysis of differentially accessible peaks was performed using FindAllMarkers(). ChromVAR (Schep et al., 2017) was used to analyze differential transcription factor binding motif activities between groups of cells. The expression features of marker genes and the projection of motif enrichment scores was profiled and visualized using the R package ggplot2 (v.3.5.1).

### Flow cytometry analysis

Single cell suspensions from dissociated lungs were prepared the same as described in the method of cell sorting. Then cells were stained with a viability dye (Ghost Dye Red 780, TONBO), followed by incubation with Fc blocking antibody (BD). Cells were stained with fluorochrome-conjugated monoclonal antibodies (Biolegend) against the following surface markers: AF488-conjugated anti-CD11b, Alexa Fluor 700-conjugated anti-F4/80, BV510-conjugated anti-CD45, BV605-conjugated anti-SiglecF, V450-conjugated anti-TCRβ, PE-conjugated anti-MHC II, APC-conjugated anti-CD11c, PerCP-Cy5.5-conjugated anti-Ly6C and PE-Cy7-conjugated anti-Ly6G. Flow cytometry was performed on an LSR II (BD) at Flow Cytometry Core in VA San Diego Health Care System and La Jolla Institute for Immunology. FlowJo™ v10.8 Software (BD Life Sciences) was used for data analysis.

### Data availability

Raw and processed files from scRNA-seq and scATAC-seq have been uploaded to the GEO database under accession number GSE328025.

## SUPPLEMENTAL INFORMATION

Figures S1–S5, Videos 1 and 2, Tables S1 and S2

## ACKNOWLEDGEMENTS

The authors thank Sun lab members for discussions. UCSD Microscopy Core supported by NINDS-P30NS047101. NovaSeq 6000 in IGM core was purchased using NIH S10-OD026929. This work was supported by NIH 1R01HL172027-01 and 1R01HL169853-01 (to X.S.) and NIH/NHLBI Pathway to Independence K99/R00HL171830 (to L.X.).

## AUTHOR CONTRIBUTIONS

L.X. and X.S. conceived and designed experiments. L.X., K.H.C., B.D., B.P.Y., R.B.L., X.G.W., Z.H.Z., V.Y., R.S. and P.S.B. performed experiments. L.X., K.H.C., B.D., B.P.Y. and M.Z.G. analyzed data. X.G.W., Z.H.Z. and M.B. procured mouse CLAD samples. Z.B., D.N.K., O.E. and M.B. provided advice. L.X. and X.S. wrote the manuscript.

---

**Figure S1.**
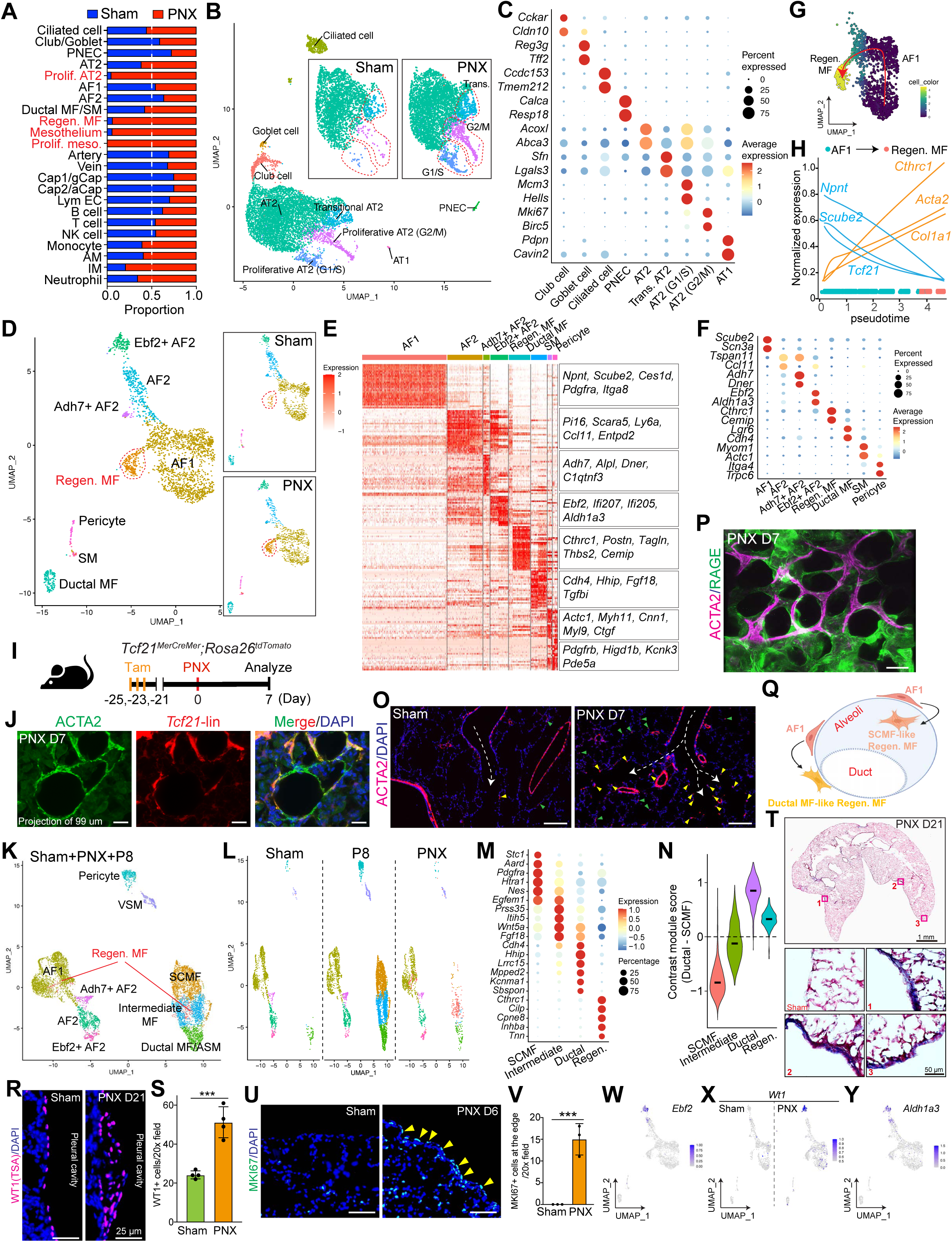
Characterization of epithelial and mesenchymal cells during lung regeneration post PNX. Related to Figure 1. (A) Ratio of normalized cell numbers from sham and PNX lungs for each cell type. (B) UMAP embedding of integrated epithelial cells from sham and PNX lungs at day 7 post surgery. AT2-related cells are separated into sham and PNX conditions in the boxes. (C) Dot plot profiling of marker genes of each cell type. (D) UMAP embedding of integrated mesenchymal cells from sham and PNX lungs at day 7 post surgery. (E-F) Profiling of marker genes in each mesenchymal cell type, as presented by the heatmap. Representative genes were listed in the box at right. Dot plots of selected genes were shown in (F). (G) Differentiation trajectory of regenerative MF, as presented by pseudotime analysis from Monocle 3. (H) Expression changes of representative genes along the differentiation pseudotime. (I) Schematic of experimental procedure used to assay cellular origin of regenerative MF post PNX in *Tcf21MerCreMer;tdT*omato mice. (J) Representative images of ACTA2 staining with lineage marker tdTomato at day 7 post PNX. Scale bars, 20 µm. (K-L) UMAP embedding of integrated mesenchymal cells from sham, PNX and postnatal day 8 lungs. Cells from different conditions were separately shown in (L). (M) Dot plot profiling of marker genes of each myofibroblast-featured cell type. (N) Quantification of SCMF- or Ductal MF-biased cell states in each cell type, as presented by contrast module scores of SCMF- and Ductal MF-related gene sets. (O) Immunostaining of ACTA2 in sham and PNX lungs at day 7 post surgery. Arrowheads denote regenerative MFs in either the ductal (yellow) or alveolar region (green). Airways were denoted by dashed white lines. Scale bars, 20 µm. (P) Maximum intensity projection of ACTA2 and AGER staining at day 7 post PNX. Scale bars, 50 µm. (Q) Schematic summary of cellular origins of regenerative MF. (R-S) Representative immunofluorescent labeling of WT1 at the lung pleura from sham and PNX at 7 days post surgery. Quantifications were shown in (S). n=4 for each group. Scale bars, 20 µm. (T) Trichrome staining in the lung at day 21 post PNX. (U-V) Representative immunofluorescent labeling of MKI67 at the lung pleura from sham and PNX at 7 days post surgery. Quantifications were shown in (V). n=3 for each group. Scale bars, 20 µm. (W) Profile of *Ebf2* expression on the integrated UMAP of mesenchymal cells, as presented by feature plot. (X) Profile of *Wt1* expression on the UMAPs of mesenchymal cells separated by sham and PNX, as presented by feature plot. (Y) Profile of *Aldh1a3* expression on the integrated UMAP of mesenchymal cells, as presented by feature plot.

**Figure S2.**
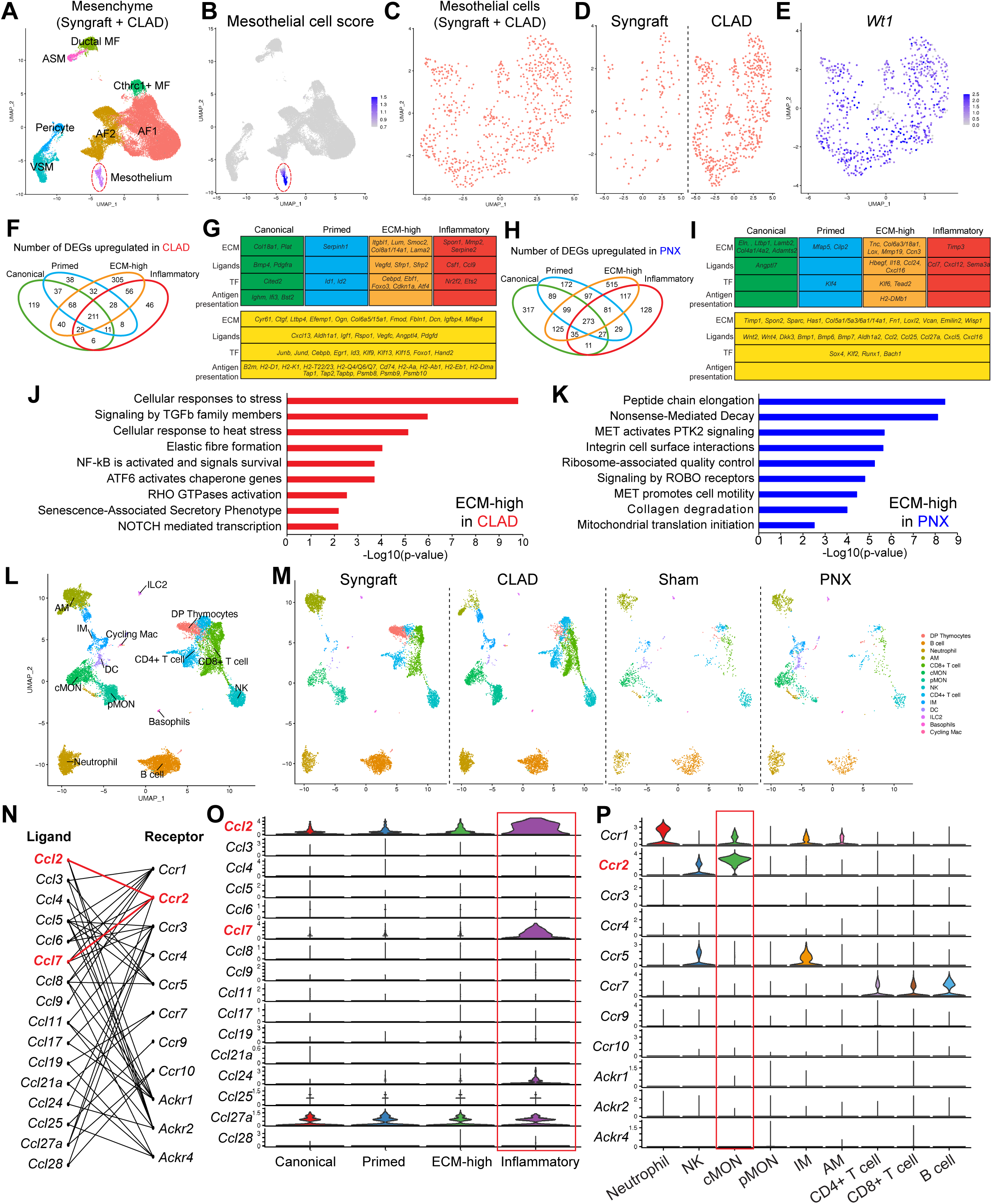
PNX-induced inflammatory mesothelial cells crosstalk with myeloid cells. Related to Figure 2. (A) UMAP embedding of integrated mesenchymal cells from syngraft and CLAD lungs. Mesothelial cells are highlighted by the red dashed circle. (B) Profile of module score of mesothelial cells on the integrated UMAP by feature plot. (C-D) UMAP-based reclustering of integrated mesothelial cells from syngraft and CLAD lungs. Cells from individual conditions were separately shown in (D). (E) Profile of *Wt1* expression on the integrated UMAP by feature plot. (F, H) Venn diagrams showing the number of upregulated genes uniquely present or shared in each of the subtypes of mesothelial cells in CLAD (F) or PNX (H). (G, I) Representative genes uniquely present or shared in each of the subtypes of mesothelial cells in CLAD (G) or PNX (I). DEGs shared by at least two subtypes are listed in the yellow boxes. (J-K) Gene ontology analysis of DEGs in ECM-high mesothelial cells between CLAD in (J) versus PNX in (K). (L-M) UMAP embeddings of integrated immune cells from syngraft, CLAD, sham and PNX lungs. Cells from different conditions were separately shown in (M). (N) Ligand-receptor pairs in the CCL family that are expressed among interactions between inflammatory mesothelial cells and immune cells. (O-P) Gene expression profiling of CCL2 ligands in expanded mesothelial cells (O) and receptors in immune cells (P), as presented by violin plots.

**Figure S3.**
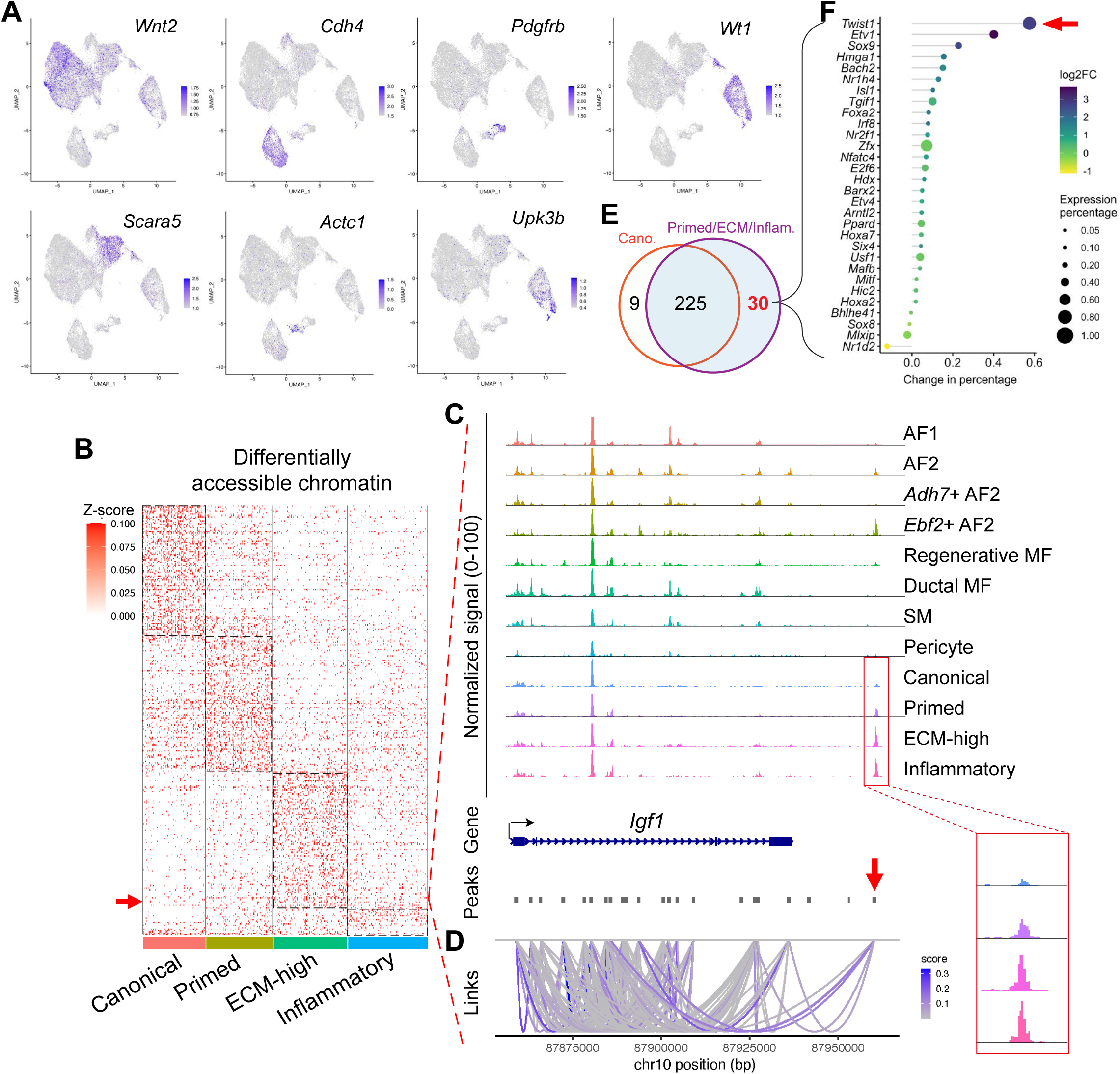
Characterization of scATAC-seq. Related to Figure 3. (A) Expression profiling of marker genes of each cell type, as presented by feature plots. (B) Differentially accessible chromatin of each cell subgroup in the expanded mesothelial cells. (C-D) Chromatin accessibility dynamics of *Igf1* in the mesenchymal cells, as presented by coverage plots. Cis-regulatory DNA element interactions predicted by Cicero were shown (D). (E) Number of enriched TFs from the predicted GRNs in canonical and activated mesothelial cells. (F) A ranking of TFs in (E), which are exclusively enriched in activated mesothelial cells, based on their relative gene expression changes. Values of x-axis are calculated by subtracting gene expression percentage in canonical mesothelial cells from the gene’s percentage in activated mesothelial cells.

**Figure S4.**
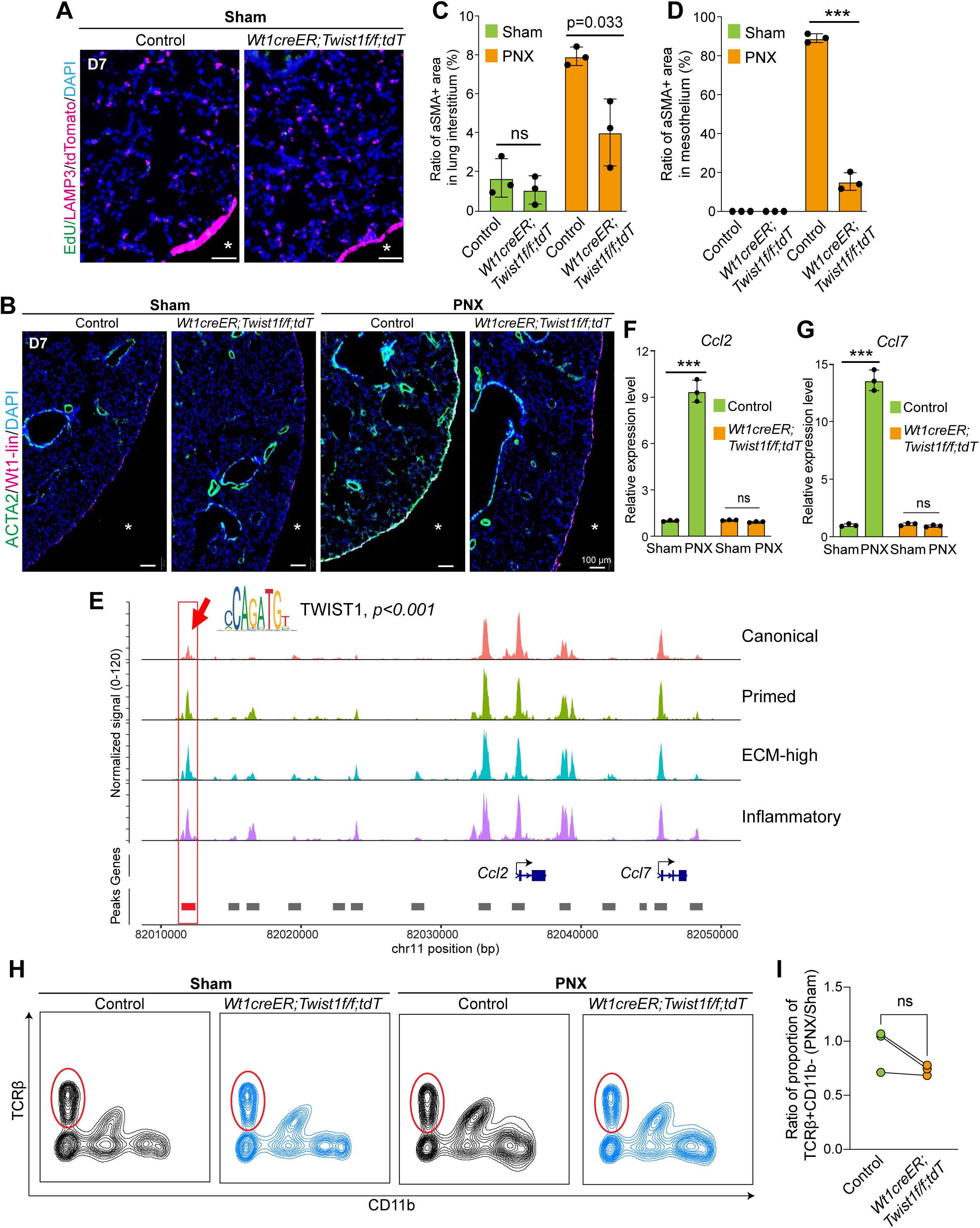
TWIST1 is required for mesothelial expansion. Related to Figure 4. (A) Representative immunofluorescent labeling of EdU signals on top of LAMP3 in the lung of *Wt1creER;Twist1;tdT* mutants and relative controls at 7 days post sham. Scale bars, 20 µm. (B-D) Representative immunofluorescent labeling of ACTA2 with *Wt1* lineage marker tdTomato from the lung of *Wt1creER;Twist1;tdT* mutants and relative controls at day 7 post surgery. Quantifications of ACTA2 staining in the interstitium and pleura were shown in (C) and (D), respectively. n=3 for each group. Scale bars, 20 µm. (E) Track view of ATAC-seq signal around *Ccl2* and *Ccl7*. The boxed region highlights the DNA regulatory element predicted to be bound by TWIST1 (JASPAR motif MA1123.2). (F-G) Quantification of *Ccl2* (F) and *Ccl7* (G) in the lung of *Wt1creER;Twist1;tdT* mutants and relative controls at day 7 post surgery, as assayed by RT-qPCR. n=3 for each group. (H-I) Flow cytometry analysis of TCRβ+CD11b− T cells as gated in live CD45+ singlet in the lung of *Wt1creER;Twist1;tdT* mutants and littermate controls at 7 days post surgery. Quantifications within littermates were shown in (I). n=3 for each group.

**Figure S5.**
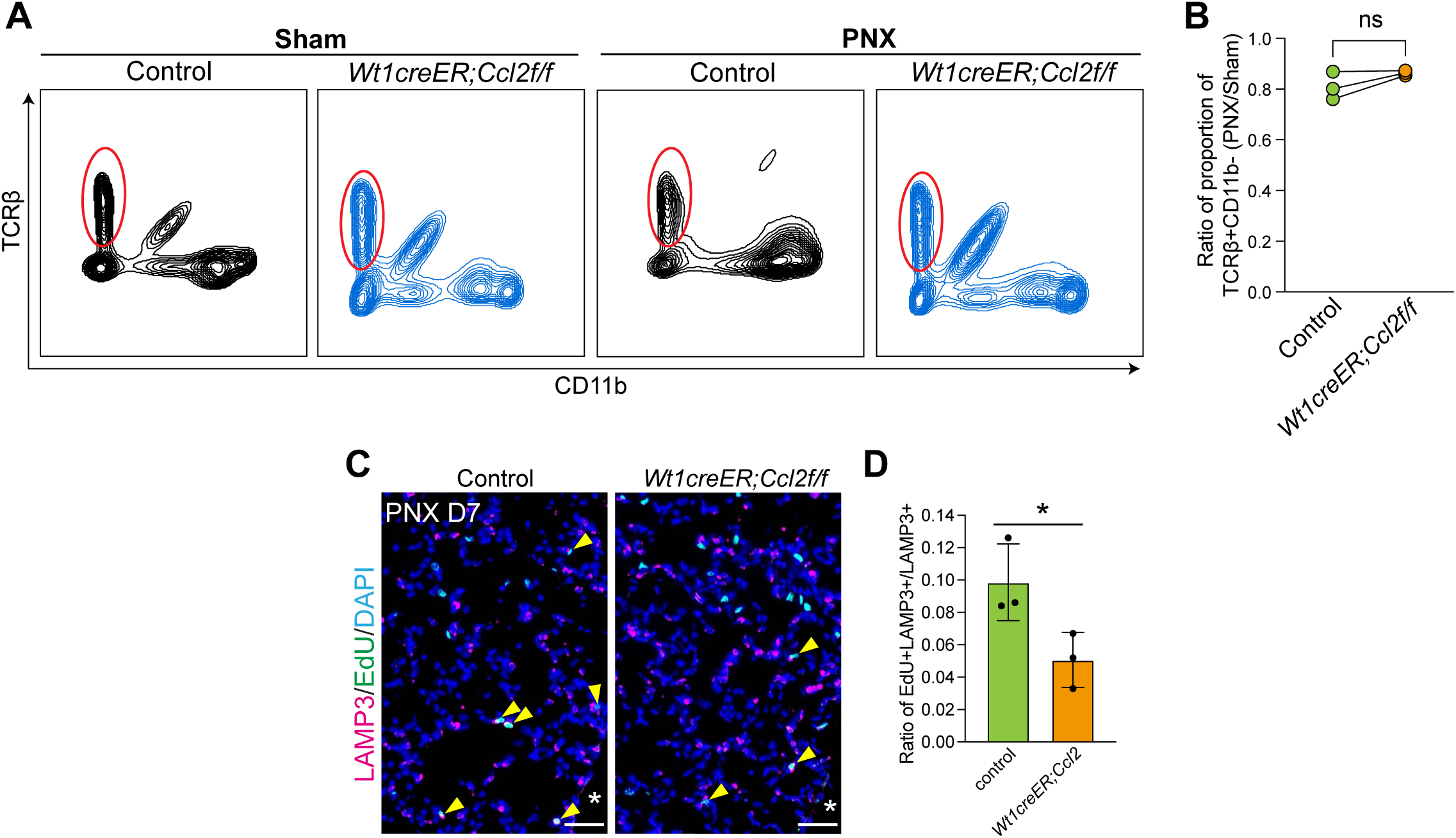
Mesothelial cell-derived CCL2 is required for AT2 proliferation post PNX. Related to Figure 5. (A-B) Flow cytometry analysis of TCRβ+CD11b− T cells as gated in live CD45+ singlet in the lung of *Wt1creER;Ccl2* mutants and littermate controls at 7 days post surgery. Quantifications within littermates were shown in (B). n=3 for each group. (C-D) Representative immunofluorescent labeling of EdU signals on top of LAMP3 in the lung of *Wt1creER;Ccl2* mutants and relative controls at day 7 post PNX. Quantifications were shown in (D). n=3 for each group. Scale bars, 20 µm.

**Table S1.**
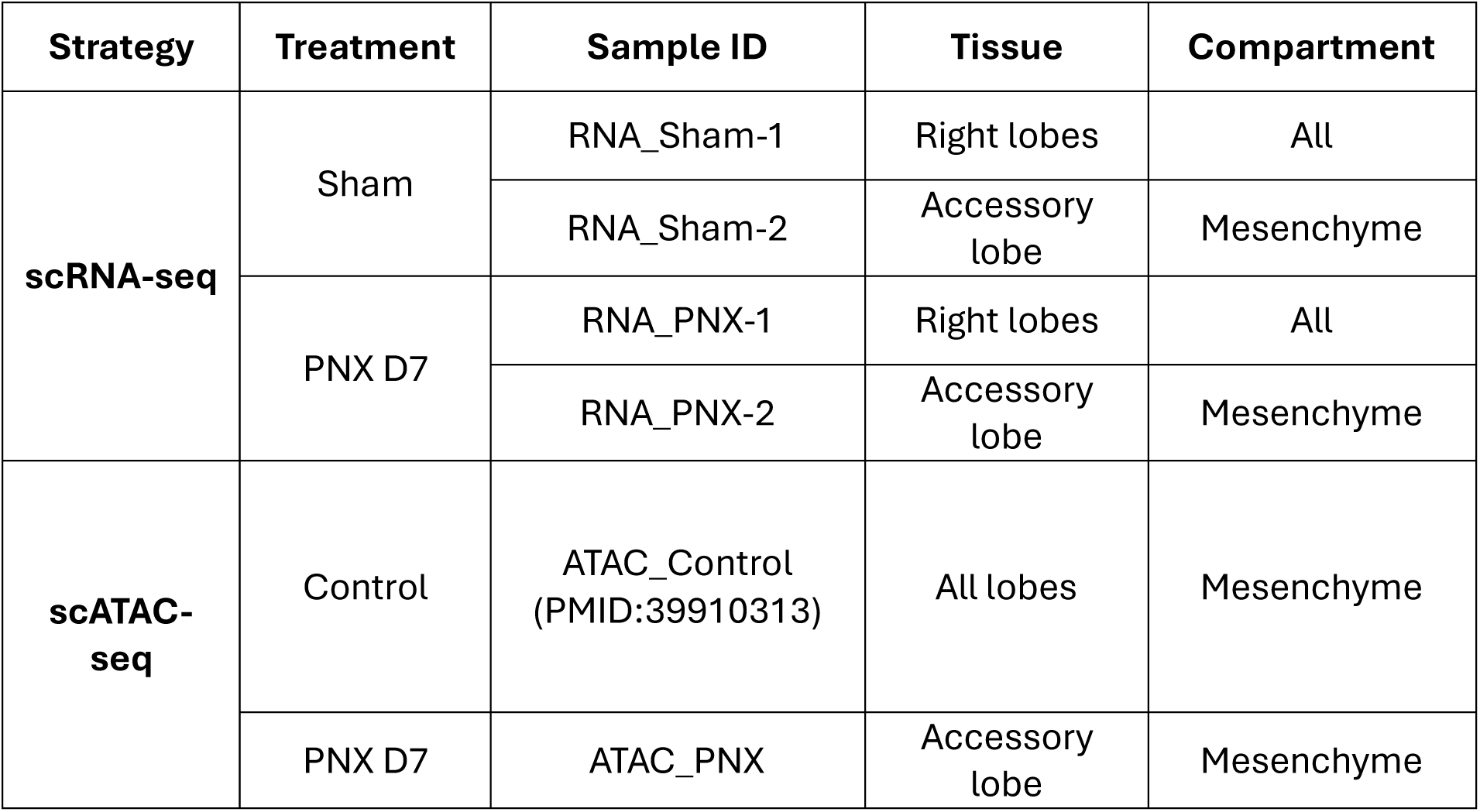
Lung samples of Sham/PNX for scRNA-seq and scATAC-seq. Related to Figures 1-5, S1-S and S4.

**Table S2.**
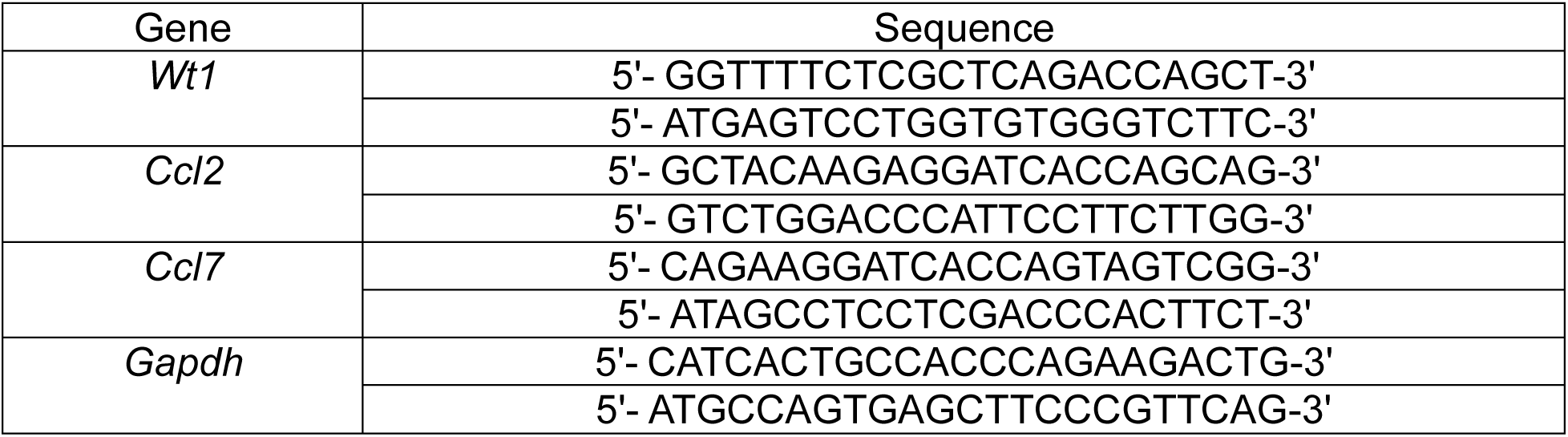
Primer sequences used for qPCR. Related to Figures 1, 4, 5 and S4.

